# Local Connectivity and Synaptic Dynamics in Mouse and Human Neocortex

**DOI:** 10.1101/2021.03.31.437553

**Authors:** Luke Campagnola, Stephanie C Seeman, Thomas Chartrand, Lisa Kim, Alex Hoggarth, Clare Gamlin, Shinya Ito, Jessica Trinh, Pasha Davoudian, Cristina Radaelli, Mean-Hwan Kim, Travis Hage, Thomas Braun, Lauren Alfiler, Juia Andrade, Phillip Bohn, Rachel Dalley, Alex Henry, Sara Kebede, Alice Mukora, David Sandman, Grace Williams, Rachael Larsen, Corinne Teeter, Tanya L. Daigle, Kyla Berry, Nadia Dotson, Rachel Enstrom, Melissa Gorham, Madie Hupp, Samuel Dingman Lee, Kiet Ngo, Rusty Nicovich, Lydia Potekhina, Shea Ransford, Amanda Gary, Jeff Goldy, Delissa McMillen, Trangthanh Pham, Michael Tieu, La’Akea Siverts, Miranda Walker, Colin Farrell, Martin Schroedter, Cliff Slaughterbeck, Charles Cobb, Richard Ellenbogen, Ryder P Gwinn, C. Dirk Keene, Andrew L Ko, Jeffrey G Ojemann, Daniel L Silbergeld, Daniel Carey, Tamara Casper, Kirsten Crichton, Michael Clark, Nick Dee, Lauren Ellingwood, Jessica Gloe, Matthew Kroll, Josef Sulc, Herman Tung, Katherine Wadhwani, Krissy Brouner, Tom Egdorf, Michelle Maxwell, Medea McGraw, Christina Alice Pom, Augustin Ruiz, Jasmine Bomben, David Feng, Nika Hejazinia, Shu Shi, Aaron Szafer, Wayne Wakeman, John Phillips, Amy Bernard, Luke Esposito, Florence D D’Orazi, Susan Sunkin, Kimberly Smith, Bosiljka Tasic, Anton Arkhipov, Staci Sorensen, Ed Lein, Christof Koch, Gabe Murphy, Hongkui Zeng, Tim Jarsky

**Affiliations:** Allen Institute for Brain Science, Seattle, WA, USA; Byte Physics, Berlin, Germany; The Ben and Catherine Ivy Center for Advanced Brain Tumor Treatment, Swedish Neuroscience Institute, Seattle, WA, USA; Department of Neurological Surgery, University of Washington, Seattle, WA, USA; Epilepsy Surgery and Functional Neurosurgery, Swedish Neuroscience Institute, Seattle, WA, USA; Department of Pathology, University of Washington, Seattle, WA, USA; Regional Epilepsy Center at Harborview Medical Center, Seattle, WA, USA

## Abstract

To elucidate cortical microcircuit structure and synaptic properties we present a unique, extensive, and public synaptic physiology dataset and analysis platform. Through its application, we reveal principles that relate cell type to synapse properties and intralaminar circuit organization in the mouse and human cortex. The dynamics of excitatory synapses align with the postsynaptic cell subclass, whereas inhibitory synapse dynamics partly align with presynaptic cell subclass but with considerable overlap. Despite these associations, synaptic properties are heterogeneous in most subclass to subclass connections. The two main axes of heterogeneity are strength and variability. Cell subclasses divide along the variability axis, while the strength axis accounts for significant heterogeneity within the subclass. In human cortex, excitatory to excitatory synapse dynamics are distinct from those in mouse and short-term plasticity varies with depth across layers 2 and 3. With a novel connectivity analysis that enables fair comparisons between circuit elements, we find that intralaminar connection probability among cell subclasses exhibits a strong layer dependence.These and other findings combined with the analysis platform create new opportunities for the neuroscience community to advance our understanding of cortical microcircuits.

## Introduction

The study of cortical connectivity has established “canonical” circuit diagrams among cells defined by their most accessible properties ^(1–3)^. More recently, cell subclasses defined by their long range projections and molecular markers have facilitated more detailed cortical microcircuit representations ^(4–9)^. Transcriptomic delineation of cell types currently offers the most refined description ^(10–13)^, however, tools that target transcriptomic cell types are not available. Though our knowledge of cell types has advanced, a complete description of the connectivity and synaptic properties among cell subclasses in each cortical layer is still lacking ^(14)^.

Cortical synapses are dynamic, varying their response strength in ways that are highly stochastic and also modulated by the prior history of activity. Ongoing activity affects both the probability of vesicle release and the quantal response amplitude, causing a combination of short-term facilitation and depression (transient strengthening or weakening of the synaptic “weight”). The dynamical properties of cortical synapses are influenced by both the presynaptic and postsynaptic cell types ^(15–20)^ and endow neuronal networks with essential sources of computational diversity ^(21, 22)^. Synaptic physiology studies rarely include comprehensive descriptions of these dynamics, despite their functional importance.

The mouse is a useful model for studying cortical microcircuitry because it affords a high degree of genetic and experimental accessibility. However, if we wish to understand our cognitive abilities, it is necessary to clarify unique features of the human cortical microcircuit ^(23)^. Access to postoperative tissue specimens has led to the identification of several differences between the mouse and human cortex, in particular differences among excitatory cells ^(24)^ and synapses ^(25)^. Human supragranular cortex appears to have more excitatory cell types, some of which have intrinsic properties that vary with cortical depth ^(24)^. We include experiments from human tissue in our survey in order to further identify unique and conserved features of the human cortical microcircuit as it relates to cell subclass.

Using new analyses and models, we have characterized the connectivity and properties of over 1000 synapses, offering a more comprehensive view of the diversity of cortical synapse types. We include analyses of the strength and probability of chemical and electrical connections; the latency and kinetic properties of synaptic responses in both voltage- and current-clamp; modeling of quantal release and short-term plasticity; and cell classification features such as morphology, intrinsic physiology, transgenic markers, and cortical layer. In our analysis we have given extra attention to generate measurements that are appropriate for biophysical modeling and that can be more readily compared to future studies, which has not been done by previous large-scale synaptic physiology studies ^(4, 6, 7, 19)^. We leverage these data and tools to ask how synaptic properties vary with cell type and find that excitatory dynamics align with target cell subclass, whereas inhibitory dynamics follow different rules depending on the subclass.

Synaptic variability is a primary driver of these cross-subclass differences, and also distinguishes human excitatory synapses which have dramatically lower variability. We also find that a single circuit representation fails to capture the diversity of intralaminar connectivity among cell subclasses in different layers. All data and tools are available from our web portal at https://portal.brain-map.org/explore/connectivity/synaptic-physiology.

## Results

We performed 1,853 experiments (Fig 1A) in acute brain slices targeting Layer 2 (L2) to L6 of young adult mouse primary visual cortex (VISp; 1,645 experiments) and human fronto-temporal cortex from neurosurgical excised tissue (208 experiments). We utilized transgenic mice that express unique reporters in two subclasses ^(26)^ (Table S1). Six excitatory subclasses were layer or projection-class specific (Nr5a1 and Rorb, L4; Sim1 or mscRE4-FlpO AAV, L5 ET; Tlx3, L5 IT; Ntsr1, L6 CT) while three inhibitory subclasses (Pvalb, Sst, Vip) were assessed in all targeted layers (Fig 1A). We probed 22,833 potential connections (mouse: 20,287; human: 2,546) of which 1,665 were connected by chemical synapses, giving an overall connectivity rate of 7.3% (mouse: 1,466 (7.2%); human: 199 (7.8%)).

**Figure 1.**
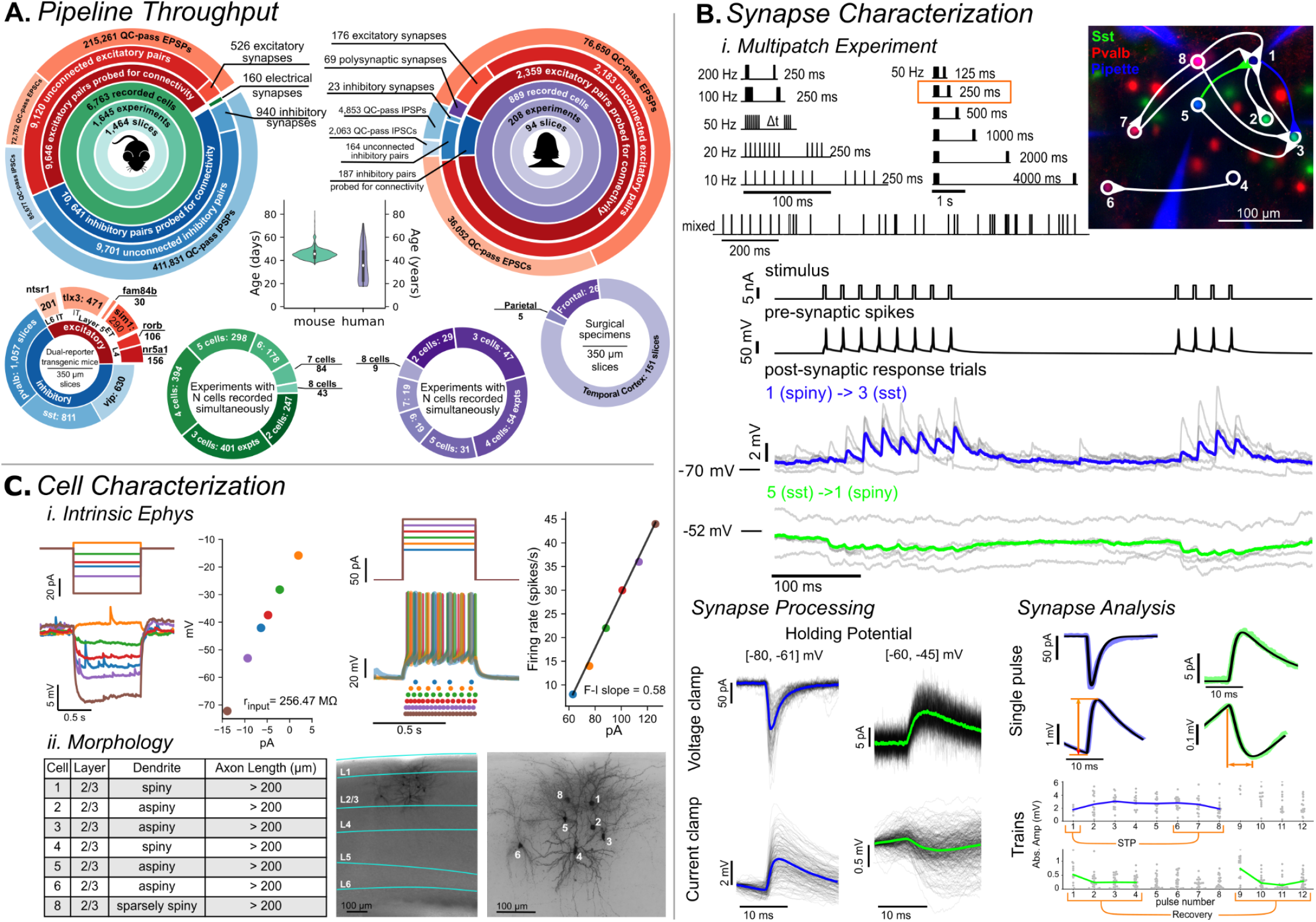
Synaptic Physiology Pipeline: **A**. Throughput of the pipeline. Each large circle shows statistics for data collected from mouse primary visual cortex (left) and human (right). The center plot shows the age distribution of each organism. Below, the outer small circles show the distributions of recorded cells; in mouse (left) this is denoted by transgenic subclass of excitatory and inhibitory cells and in human this is denoted by cortical region. The inner circles show the number of experiments in which two cells were recorded from, three cells, etc., up to a maximum of eight cells being simultaneously recorded. **B**. *i.Multipatch Experiment*: The top panel shows the set of stimuli used in a multipatch experiment. They consisted of pulse trains at various set frequencies with 8 initial pulses, a delay of 250 ms, and then four more pulses. In the 50 Hz stimulus various delays were interposed between the first 8 and second 4 pulses. The mixed frequency stimulus started with 8 pulses at 30 Hz and then proceeded with 30 pulses at set, but variable instantaneous frequencies ranging from 10-200 Hz. The fluorescent images to the right show cells recorded in an example experiment with the connectivity diagram overlaid. Overall in this experiment, cell 1, an unlabeled spiny cell, received convergent input from both Sst cells (3, 5) and Pvalb cells (7, 8), while cell 8, a Pvalb cell, showed divergent outputs to three different subclasses. Thus, ten synaptic connections were identified in this experiment, characterizing seven of the possible nine interaction types. Below is an example of the electrophysiology recording of cells 1, 3, and 5 during the 50 Hz stimulus highlighted by the orange box. The stimulus and presynaptic spikes were recorded from cell 1 which was presynaptic to cell 3. The response of cell 3 to individual presentations (trials) of the stimulus are shown in grey and the average in blue. A similar example for an inhibitory connection also found in this experiment from cell 5 to cell 1 is shown in green. *ii. Synapse Processing*: A synapse was identified by aligning every presynaptic spike and overlaying the postsynaptic response (black traces) in both voltage clamp (top row) and current clamp (bottom row) with the average in color (colors correspond to the same synapses in the *Multipatch Experiment* panel). Excitatory connections were evaluated at a holding potential between −80 and −61 (left column) mV while inhibitory connections were evaluated at a holding potential between −60 and −45 mV (right column). *iii Synapse Analysis*: When a connection was identified the average (colored) was fit (black) and metrics such as amplitude and rise time (orange lines) were extracted from the fit output. These fit parameters were used to guide fitting of individual pulses so that we could quantify changes in synapse amplitude over the course of a train. Short term plasticity (STP) was quantified as the amplitude of the first pulse subtracted from the average of pulses 6-8, normalized by the 90th percentile amplitude for the connection overall. Recovery was similarly quantified as the average of pulses 1-4 subtracted from pulses 9-12, normalized by the 90th percentile. **C**. *i Intrinsic Ephys*: Long pulse steps were applied to quantify intrinsic cell properties, and electrical synapses (Figure S4). Hyperpolarizing steps (left) delivered to an example cell probed the subthreshold I-V relationship to quantify metrics such as input resistance while depolarizing suprathreshold sweeps (right) were used to measure spiking and firing rate properties. *ii Morphology*: Cells were filled with biocytin during recording and stained. 20x images of the full slice stained with DAPI allowed identification of the cortical layers and 63x z-stack images were used to assess morphological properties such as dendritic type and axon length. Images and cells here are the same as in (B).

Details of the experiments are described in Fig. 1B/C and in the Methods. In each experiment, up to eight neurons were selected for simultaneous whole-cell patch-clamp (multipatch) recording primarily in current-clamp, with a subset of stimuli administered in voltage-clamp (Fig 1B). Stimuli were elicited in each patched neuron, in turn, while all others were recorded for evidence of a postsynaptic response. Stimulus trains of eight pulses at frequencies ranging from 10 – 200 Hz probed the degree to which individual connections displayed short-term plasticity (STP). After a delay, four additional pulses were delivered to characterize how synapses recovered from STP. Cells were stimulated with long current pulses to characterize their intrinsic physiology and later stained with biocytin to characterize their morphology (Fig 1C).

### Intralaminar connectivity in mouse VISp

#### Distance dependence of connectivity

We probed connectivity among cells up to ∼200 μm apart (Fig 2A), but could not ensure that intersomatic distances were sampled equally across different connection elements (specific pre-post combinations). In order to make reliable comparisons, we modeled the spatial profile of connectivity versus lateral somatic distance with a Gaussian ^(27)^ and estimated the peak (*p_max_*) and lateral spread of connection probability (sigma: *σ*) using maximum likelihood estimation (see Methods, Fig S2B). This analysis provides an estimate of the spatial profile of connectivity while being relatively robust to differences in the sampled intersomatic distribution. We initially classified cell pairs among the four combinations of excitatory and inhibitory cell class (E→E, E→I, I→E, I→I). These data were well fit by the Gaussian model (Fig 2A, solid red line) and indicated low peak connectivity among excitatory cells (∼5%), moderate connectivity rates among inhibitory cells (∼11%), and higher connectivity across E-I cell classes (E→I 12%, I→E 15%). In a rabies tracing study of primary visual cortex L2/3, E→E connections were found to have a wider lateral extent compared to I→E ^(28)^. We confirm this result and additionally find that E→E and I→I connections have a similar spatial extent that is larger than both E→I and I→E connections. Taken together, these results suggest a spatial wiring principle where within-class connectivity has a wider spatial profile than across-class connectivity.

**Figure 2.**
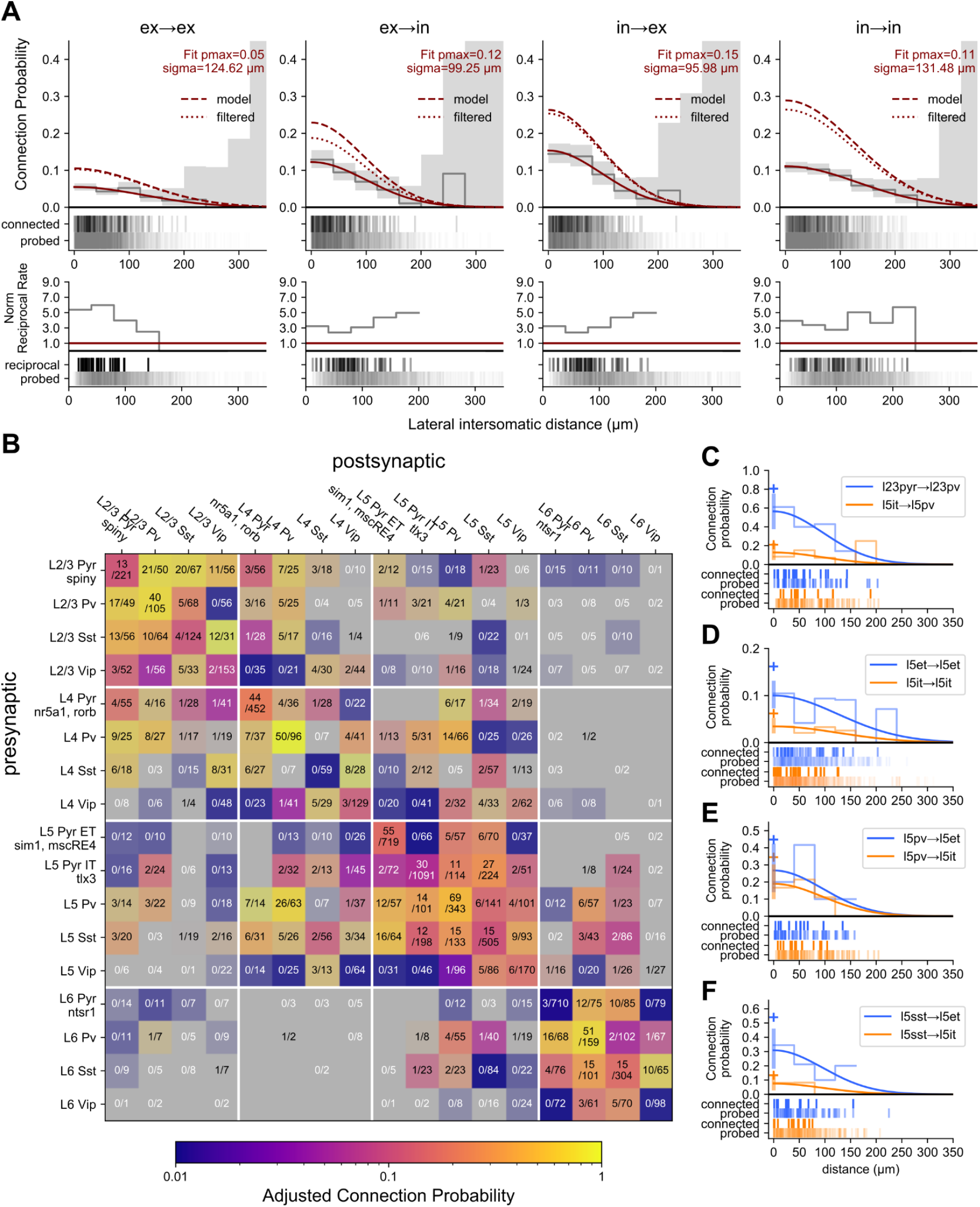
Connectivity: **A.** Gaussian fit of connection probability as a function of lateral intersomatic distance. All mouse cells were divided into two main classes, excitatory and inhibitory, and pairs classified into the four combinations of those two classes. *Top row*: Connection probability as a function of intersomatic distance was fit with a Gaussian (red line) and output parameters *p_max_* and sigma (*σ*) describe the max connection probability and width of the Gaussian. Adjusted connection probability as a function of intersomatic distance adjusted for presynaptic axon length, depth of the pair from the slice surface, and detection power of connections using a unified model (dashed red line) or via filtering of the data (dotted red line) yield similar results (see Results/Methods). Grey line and area are 40 μm binned average connection probability and 95% confidence interval. Raster below shows distance distribution of connections probed (bottom) and found (top). *Bottom row*: Normalized rate of reciprocal connections. Probed pairs are unordered and the number of reciprocal connections counted was normalized to the expected value of connection probability squared for a randomly connected network (solid red line). **B**. Connection probability matrix for mouse. Connection probability is estimated using a unified model accounting for all corrections as determined from A (dashed red line, “model”). The shading of each element indicates the 95% CI of the data with higher contrast indicating smaller CI and lower contrast (toward grey) indicating larger CI. The number of connections found out of the number of connections probed are printed in each element. Excitatory cell class in layers 4-6 were distinguished by transgenic reporters while layer 2/3 excitatory cells are defined as morphologically spiny. Three inhibitory classes Pvalb, Sst, and Vip are defined by transgenic reporters. **C-E**. Gaussian fit of connection probability vs intersomatic distance (with CI at *p_max_*, shaded region) for two contrasting elements with connections found and connections probed raster below. Cross symbol denotes *p_max_* with all adjustments.

An overabundance of bidirectional connections relative to unidirectional connections can be evidence for connectivity rules that promote the formation of bidirectional connections ^(29)^. A simpler explanation, however, is that bidirectionality is not specifically promoted in cortex, but rather an artifact of merging connectivity results across cell types ^(30)^. We quantified the ratio of connected pairs with and without bidirectional connections and observed that reciprocal connections were 3 to 5 times more common than expected for a randomly connected network (red line) among class-level connections (Fig 2A, bottom row). Further analysis of higher order connectivity motifs in our data and their impact on cortical computation is pursued in a parallel study ^(31)^.

#### Connection probability measurement; slicing artifacts and detection limits

In the *in vitro* slice preparation, some connections may be severed, reducing the measured rate of connectivity ^(27, 32)^. To mitigate this bias, we used thick slices (350 μm), targeted cells deep in the slice (Fig S1B, median cell depth = 75 μm), and focused on local (<200 µm apart), intralaminar connections. By modeling the effects of cell depth and axon length on connection probability, we were able to estimate the size of this bias and adjust our *p_max_* measurements accordingly. This yielded an ∼5-10% increase in *p_max_* when accounting for severed axons (Fig S1A, D) and a more modest ∼5-20% increase in *p_max_* when accounting for depth of the targeted cells from the slice surface (Fig S1B, D).

In addition to slicing artifacts, background electrical noise may obscure synaptic responses and reduce the observed connection probability. Conversely, increasing the number of presynaptic action potentials enhances the likelihood of detecting a connection. To account for these biases, we quantified the ‘detection power’ for each pair of cells that were probed for connectivity (see methods). A model of the relationship between detection power and connection probability resulted in a 30-50% increase in estimated *p_max_* for excitatory connections and up to 3-fold increase for inhibitory connections (Fig S1D), suggesting that the observed connectivity rate is affected more by detection power than by slicing artifacts. Detection power is seldom reported and may explain cases where the observed connectivity *in vitro* is higher than *in vivo* for the same brain area and connections ^(33, 34)^.

We extended our model of connection probability on intersomatic distance (Fig 2A, solid line) to include the effects of slicing and response detection outlined above (see Methods). The model-adjusted connectivity rate resulted in a 2-3 fold increase in estimated *p_max_* at the cell class level (Fig 2A dashed line). To confirm these results, we implemented the connectivity adjustment by filtering the data for axon lengths, pair depths, and detection power values above their respective median (Fig 2A, dotted line) where the impact on connection probability is reduced (Fig S1A-C). This yielded a comparable increase in estimated *p_max_* (Fig 2A dotted vs dashed line) and as such we used the model to adjust peak connectivity estimates at the subclass level where filtering would induce under-sampling. Adjustments for presynaptic axon length and pair depth were relatively uniform across subclasses, whereas inhibitory connections received a larger adjustment for detection power compared to excitatory connections (Fig S1E).

#### Connectivity among cell subclasses

The intralaminar connectivity among mouse VISp subclasses is summarized as a matrix in Fig 2B (S2A). The hue of each element indicates the model adjusted pmax with the saturation scaled by the 95% confience interval (more saturated colors have smaller confidence intervals, or CIs). Often, we observe connectivity results that are, across all layers, consistent with prior results such as strong recurrent Pvalb connectivity, indiscriminate inhibition from Sst cells, and Vip disinhibition of excitatory cells via Sst ^(4, 6, 7)^. Cortical disinhibition via the Vip→Sst pathway is a well described phenomenon ^(35, 36)^ however, we also see high connectivity in the reverse direction (Sst→Vip), the implications of which are less well understood.

We also saw that intralaminar connection probability varies by layer (Fig 2B and C), long-range projection target of excitatory cells (Fig 2D), and cell subclass more generally (Fig 2 E, F). Our results show that L2/3 connectivity is substantially higher than other layers, whereas L5 has overall lower connectivity. For example, the pyramidal to Pvalb connection in L2/3 (*p_max_*=0.76 [0.49, 1.00] CI) compared to L5 (IT *p_max_*=0.21 [0.11, 0.35] CI) (Fig 2C, S2A). Additionally, recurrent connections between excitatory and Vip cells are common in L2/3 (E→Vip 0.51 [0.28, 0.79]; Vip→E 0.13 [0.03, 0.31]) but rare or absent in deeper layers (Fig S2A). Conversely, Pvalb→Vip connections were found in all layers except L2/3.

Within L5 we found several differences between two excitatory projection classes, intratelencephalic (IT, labeled by Tlx3) and extratelencephalic (ET, labeled by Sim1 and mscRE4). ET cells overall have more input from local sources relative to IT cells. ET cells have higher recurrent connectivity (Fig 2D) as well as receive unidirectional input from IT cells, consistent with previous results ^(37)^. Sst cells also innervate ET cells at a higher rate than IT cells (Fig 2E-F); a similar connectivity pattern was observed in rat frontal cortex ^(38)^.

Sst is thought to avoid connecting with itself ^(4, 11, 39)^, however, we observe connections between Sst neurons in almost every layer (Fig 2B). It is known that the Sst-IRES-Cre driver can sparsely label fast-spiking interneurons that also express Pvalb ^(40)^. UMAP projection of intrinsic properties from inhibitory cre-types showed that slightly more than half of the recurrent Sst connections had both the pre- and postsynaptic cell in a cluster that was spatially distinct from Pvalb cells, suggesting that these connections came from cells that were intrinsically Sst-like (Fig S3A). We confirmed that, in at least one case, both cells in the pair had axons extending into L1 and sparsely spiny dendrites, consistent with Sst neurons (Fig S3B). We performed follow-up experiments utilizing the Patch-seq method ^(41)^ and further confirmed that cells which transcriptionally mapped to the Sst subclass do form connections with each other (3 connections found out of 47 probed, Fig S3C).

#### Electrical synapses

In addition to chemical synaptic transmission among cell subclasses, electrical connections, facilitated by gap junctions, were also found between inhibitory subclasses (Fig S4A). The likelihood of electrical synapse connectivity as a function of lateral intersomatic distance could be approximated by a Gaussian but with a narrower profile, *σ* = 77 μm (Fig S4B), compared to chemical synapses between inhibitory cells (*σ* = 131 μm, Fig 2A; p<0.001, Fig S4B). This is consistent with previous reports of electrical connections between nearby Pvalb cells which showed that the average distance between electrically coupled cells was short, 40-80 μm ^(42–44)^. L2/3 Pvalb cells showed the highest rate of electrical connections (18/106, ∼17%), while those among Vip cells (9/772, ∼1%) were the most rare, in contrast to a previous report of Vip electrical connections which were more prevalent ^(7)^. A majority of electrical synapses were found between like subclasses (148/154, 96%) and were bidirectional (134/154, 87%). The distance of the gap junction from the soma coupled with the potential for some rectification of electrical connections ^(45)^ could account for the few cases where reciprocal electrical connections were not observed. The coupling coefficient of electrically coupled pairs was comparable across subclass (Fig S4C) largely due to the lower input resistance of Pvalb cells (Fig S4D). Estimating junctional conductance revealed stronger electrical synapses between Pvalb (0.37 [0.20, 0.53] nS, median [interquartile range]) cells than either Sst (0.21 [0.17, 0.29] nS) or Vip (0.06 [0.02, 0.16] nS) (Fig S4C).

### Synaptic strength and kinetics

In addition to connectivity, synaptic properties determine the impact of a connection on the postsynaptic neuron and, ultimately, on cortical processing. The strength, latency, and kinetics (rise time and decay tau) of local synapses have been described across multiple studies for different cell types^(4, 46–50)^. Still, this study offers the first open dataset in which synapse properties may be directly compared across many cell subclasses and layers. Synaptic latency, rise time, decay tau, resting state amplitude, and near maximal (90^th^ percentile) amplitude were extracted from exponential fits to the average postsynaptic response (PSC/P) (see Methods for more detail).

At the class level, inhibitory synapses show short latencies (median=1.08 ms), slow kinetics, and relatively strong PSPs. These trends are largely driven by the subclass of the presynaptic cell and Pvalb in particular. Pvalb synapses are extremely fast, with sub-millisecond latencies (Fig 3A) highlighted by Pvalb→L6 excitatory synapses (0.97 [0.86, 1.09] ms, median [interquartile range]). Pvalb cells also elicit the largest resting state IPSP amplitudes regardless of postsynaptic cell subclass. One exception is the strength of Pvalb→L5 ET (−0.33 [−0.31, −0.80] mV), which is weaker than Pvalb→L5 IT (−0.53 [−0.21, −0.77] mV), again highlighting the dichotomy of these two projection classes. Presynaptic Sst cells stand out for having some of the slowest kinetics, independent of postsynaptic target (see Sst→Vip rise time: 7.15 [5.7, 10.0] ms, decay: 46.7 [24.41, 235.09] ms; Sst→L5 IT rise time: 6.5 [5.73, 8.55] ms, decay: 32.0 [21.64, 64.36] ms).

**Figure 3.**
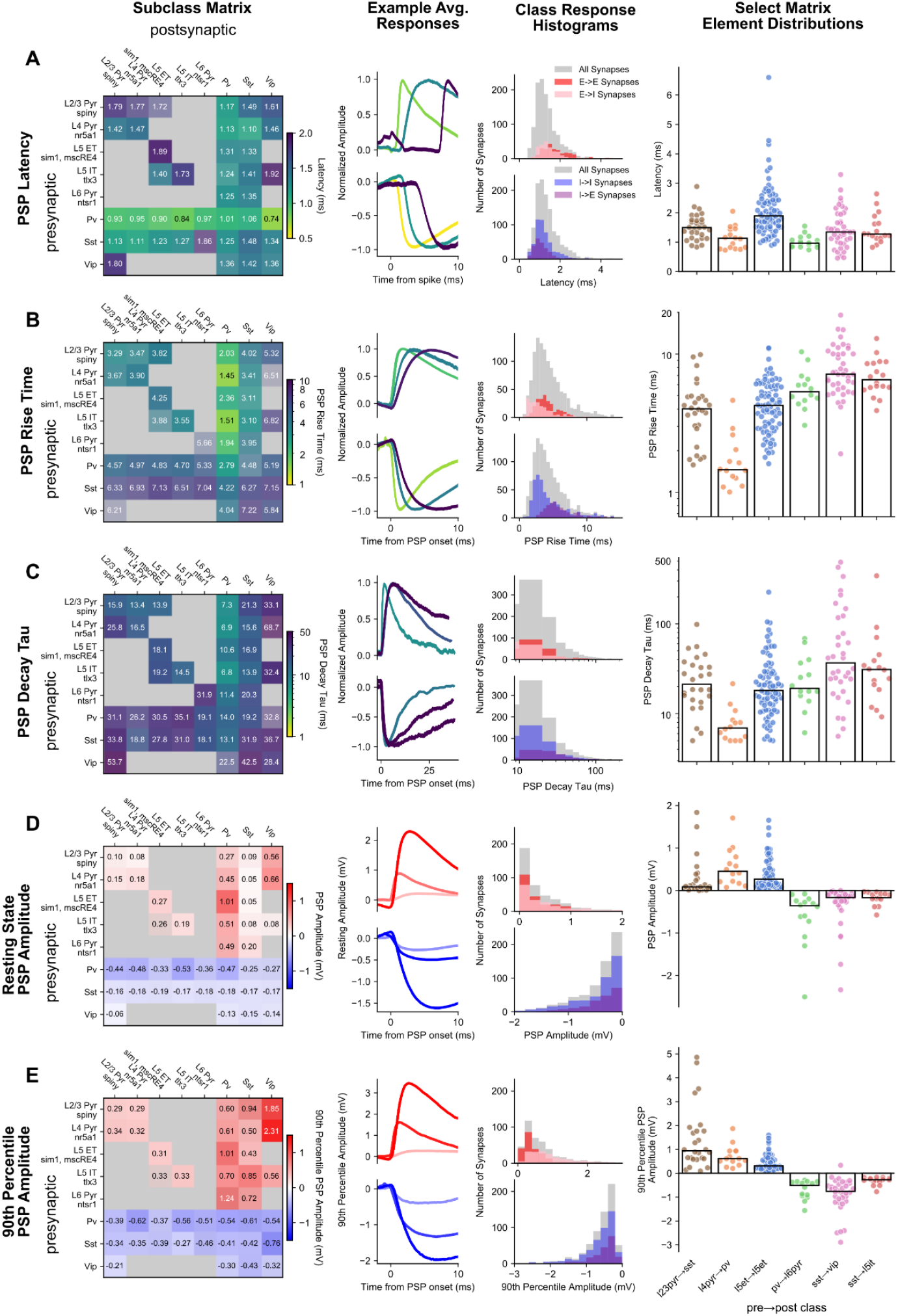
Synaptic strength and kinetics: **A.** (left to right) PSP latency matrix; excitatory and inhibitory minimum average latency, median average latency, and maximum average latency representative traces (light to dark colors); latency histograms for the major connection classes (E→E, E→I, I→I, I→E); latency summary plots for a subset of matrix elements. **B.** (left to right) PSP rise time matrix; representative traces for excitatory and inhibitory minimum average rise time, median average rise time, and maximum average rise time (light to dark colors); rise time histograms for the major connection classes; rise time summary plots for a subset of matrix elements. **C.** (left to right) PSP decay tau matrix; representative traces for excitatory and inhibitory minimum average decay tau, median average decay tau, and maximum average decay tau (light to dark colors); decay tau histograms for the major connection classes; decay tau summary plots for a subset of matrix elements. **D.** (left to right) PSP amplitude matrix; representative traces for excitatory and inhibitory minimum average amplitude, median average amplitude, and maximum average amplitude (light to dark colors); amplitude histograms for the major connection classes; amplitude summary plots for a subset of matrix elements. **E.** (left to right) PSP 90th percentile amplitude; representative traces for excitatory and inhibitory minimum average 90th percentile amplitude, median average 90th percentile amplitude, and maximum average 90th percentile amplitude (light to dark colors); 90th percentile amplitude histograms for the major connection classes; 90th percentile amplitude summary plots for a subset of matrix elements. In all matrices, inhibitory cells are merged across layers. All matrices are colorized by the median (text in each element) with the saturation scaled by the standard error.

In contrast to inhibitory synapses, excitatory synapses generally have a long latency (median=1.49 ms), fast kinetics, and weak PSPs, all of which relate more to the identity of the postsynaptic cell. Excitatory→Inhibitory synapses display faster rise times than recurrent excitatory synapses (E→I 2.73 ms, E→E 3.87 ms, ks=5.18e-12, Fig 3B). Consistent with our previous work^(51)^, recurrent excitatory connections show some of the smallest amplitudes in the resting state (eg. L5 ET→L5 ET, 0.27 [0.13, 0.5] mV) whereas E→I synapses are stronger (Fig 3D, E) and generated the single biggest PSP (15.03 mV, L5 IT→L5 Sst). E→I synapse properties can be further refined by postsynaptic cell subclass. Synapses with postsynaptic Pvalb cells (see L4 Pyr→Pvalb, Fig3) have faster kinetics (rise time: 1.45 [1.28, 1.94] ms, decay tau: 6.92 [5.4, 8.36] ms) than postsynaptic Sst cells (see L2/3 Pyr→Sst, Fig 3; rise time: 4.06 [2.99, 4.91] ms, decay tau: 25.18 [15.37, 38.88] ms). A dichotomy between Pvalb and Sst is also apparent in the resting state amplitude with larger EPSPs to Pvalb cells than Sst cells; however, the strongest excitatory connections are onto Vip cells found predominantly in superficial layers (L2/3 pyr→Vip 0.56 [0.23, 0.84] mV). Although resting state excitation was weakest on to Sst cells, resting state PSP amplitude is an underestimate of the potential impact on the postsynaptic cell particularly for synapses that strongly facilitate. When we compare the 90^th^ percentile amplitude (Fig 3E), which measures near-maximal strength, E→Sst synapses are comparatively stronger, and even surpass E→Pvalb amplitudes in some cases (see L2/3 Pyr→Sst vs L4 Pyr→Pvalb). Facilitation onto inhibitory cells further contributes to the longer tails of 90th percentile amplitudes and the rightward shift of E→I (0.74 mV) amplitudes compared to E→E (0.33 mV, ks=1.11e^−16^; Fig 3E, histograms).

A recent survey of cortical connectivity found that synapse strength positively correlated with connection probability ^(6)^. This result suggests an interesting principle of connectivity, but may also result from the reduced detectability of weaker synapses. When we assessed this relationship we found that adjusted connection probability of excitatory synapses was not correlated with synaptic strength (weighted Huber regression *r*^2^ = 0.2, *p* = 0.5) while inhibitory synapses showed a small correlation (weighted Huber regression *r^2^* = 0.4, *p* < 0.01). Recall that we see a smaller effect of detection power on excitatory compared to inhibitory connections (Fig S1C) suggesting that when detection power is sufficient, connection probability is independent of synaptic strength.

### Synaptic dynamics

The strength and kinetic properties described above characterize the synapse in response to a single presynaptic spike; however, synapses are highly dynamic. PSP amplitude evolves in predictable ways over the course of milliseconds to seconds due to STP, while also being highly stochastic from one response to the next, quantified by the coefficient of variance. Overall, synaptic dynamics follows a similar pattern to synaptic strength, wherein excitatory synapses are most strongly differentiated by the postsynaptic subclass and inhibitory synapses are differentiated by the presynaptic subclass. The STP of a synapse may result in a transient increase (facilitation), decrease (depression), or no change (pseudo-linear) in PSP amplitude over the course of a stimulus train as is seen in our data (Fig 4A) and has been described previously^(15, 20, 52–54)^. The time course of recovery from STP is an equally important property of synapses, yet one that is not well described. The variable delays we imposed between the induction and recovery pulses (Fig 1B, *Multipatch Experiment*) of our 50 Hz stimulus show that at our earliest time point (125 ms), synapses are still in their STP induced state, but that by four seconds they are largely recovered.

**Figure 4.**
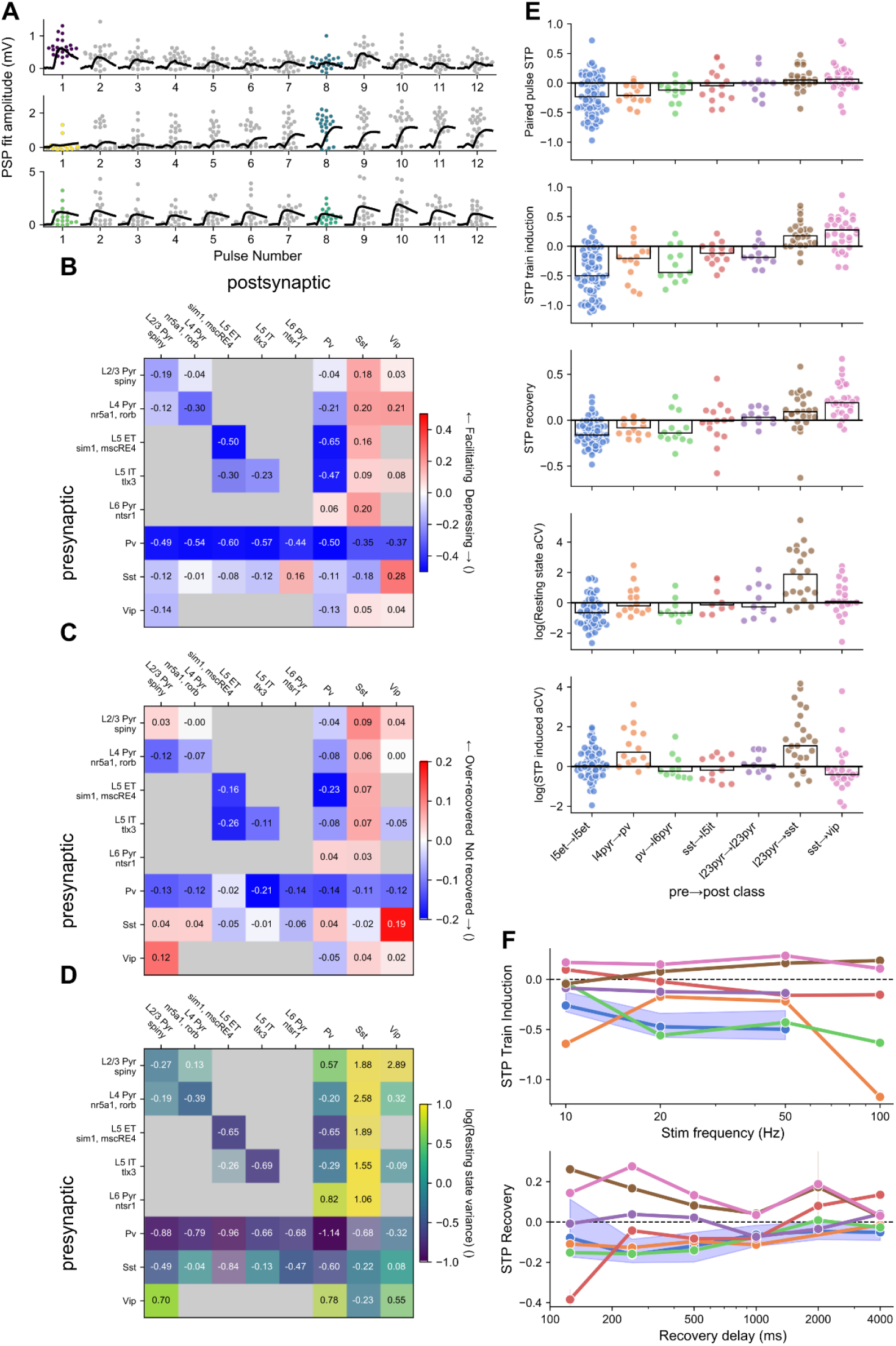
Synaptic dynamics: **A.** Representative depressing, facilitating, and pseudo-linear excitatory synapses (top to bottom) in 50 Hz train; grey/colored dots: individual PSP amplitudes; black traces: average PSP per pulse. Scatter points for pulse 1 (resting state aCV) and pulse 8 (STP induced aCV) are colored according to the color scale in **D**. **B.** Short-term plasticity matrix. **C.** Recovery (at 250ms) matrix. **D.** Resting state variance (adjusted coefficient of variation) matrix. All matrices are colorized by the median (text in each element) with the saturation scaled by the standard error. **E.** Summary plots for paired pulse ratio, STP induction ratio (avg 1st pulse amp : avg of 6th-8th pulse amp) normalized by the 90th percentile, Resting state variance, induced state variance (top to bottom). Each dot corresponds to the average response from one synapse. **F.** Train induced STP (top) at four different frequencies (10, 20, 50, 100 Hz) for each of the elements in E (colors maintained). Each dot is the grand average of all synapses in the element. For L5 ET→L5 ET the blue shading highlights the 95% confidence interval as an example. Lower plot shows recovery from STP at six different delays (125, 250, 500, 1000, 2000, 4000 ms) in a similar manner to the plot above.

Excitatory dynamics are strongly aligned with postsynaptic cell class^(54)^, and further refined by layer in the case of excitatory targets and by subclass of inhibitory targets. Recurrent excitatory connections largely depress (Fig 4B, E L5ET→L5ET), consistent with our recent study^(51)^, and show increasing depression with stimulus frequency (Fig 4F). Recurrent excitatory synapses occupy a range of recovery and variability profiles that vary with layer. Superficial layers (eg. L2/3→L2/3) tend to recover more quickly (Fig 4C) and show a higher degree of variability (Fig 4D). E→I dynamics depend on the subclass of the postsynaptic target. Excitatory to Sst cells are strongly facilitating, consistent with previous reports^(53)^. E→Sst synapses are also highly variable in the resting state, likely owing to a high initial failure rate (Fig 4A, middle, pulse 1, Fig 4E); however, strongly facilitating synapses often become more reliable in the induced state (Fig 4A, middle, pulse 8, Fig 4E). We further observe a difference in the magnitude of facilitation of synapses onto Sst cells from ET and IT cells in L5 where ET to Sst shows stronger facilitation (Fig 4B, ET→Sst 0.16 [0.12, 0.19], IT→Sst 0.09 [0.03, 0.13]). Excitatory connections onto Pvalb (Fig 4E, L4 pyr→Pvalb) were largely depressing on average, though a subset of synapses in L2/3 showed pseudo-linear STP similar to *in vivo* measurements in somatosensory cortex^(50)^. While these patterns of excitatory dynamics are apparent on average, multiple measurements of synaptic dynamics show high heterogeneity from pair to pair within a synapse type (Fig 4E).

Dynamics of inhibitory synapses show patterns more related to the subclass identity of the inhibitory presynaptic cell. Pvalb connections onto other subclasses, both excitatory (Fig 4E, Pvalb→L6 pyr) and inhibitory, are strongly depressing (Fig 4B) and still depressed at our earliest recovery time point (Fig 4C). Depressing Pvalb synapses show high reliability at the beginning of a stimulus train (Fig 4D) and become more variable in their STP induced state (Fig 4E). Connections from Sst and Vip cells skew toward depression (Fig 4E, Sst→L5 IT) but were not as depressed as Pvalb connections^(54)^. These synapses were also faster to recover, particularly Vip synapses which tended to over-recover at the shortest interval (Fig 4C).

Synaptic interactions between Sst and Vip are an exception to the trends highlighted above. Whereas most inhibitory synapses are depressing, Sst→Vip^(7)^ showed the highest degree of facilitation in our dataset (0.27 [0.06, 0.45]). The reciprocal Vip→Sst synapse is weakly facilitating, as are recurrent Vip connections. These three synapse types also over-recover on short time scales (Fig 4C) and take many seconds to fully recover (Fig 4F). Given the facilitating nature of these synapses it is interesting to note that they have only a moderate degree of variance (Fig 4D) compared to other facilitating synapses such as E→Sst.

### Human intralaminar connectivity

As a complement to the mouse visual cortex, our dataset includes synaptic physiology from human temporal cortex. Although our sampling of human synapses covered all cortical layers, our analysis focuses on the supragranular layers, which are dramatically expanded in anthropoid primate cortex ^(55)^. Previous work has shown that deep L3 cells have distinct electrophysiology, morphology, and gene expression (including genes involved in connectivity and synaptic signalling) and that many of these properties vary continuously with depth between L2 and L3b ^(24)^. Dense sampling of L2/3 allowed us to define L2, L3a, and L3b pyramidal subclasses and demonstrate that these principles of cellular diversity have correlates in synaptic physiology. These subclasses show distinct synaptic properties including unique polysynaptic connections from L2 cells and STP that closely follows the continuous variability between L2/3 subclasses.

Distance dependence of connections was modeled and adjusted as with mouse synapses, but without distinguishing connections by cell class (most synapses were recurrent excitatory). Connection probability was estimated to fall off with distance at a lateral spread (*σ*) of 140 μm (Gaussian model fit, Fig. S7A), moderately larger than the comparable value in mouse (125 μm for within-class connections), reflecting that while cortical expansion is accompanied by the scaling of neuronal morphology, much of this scaling is axial rather than lateral. Examining the connectivity between subclasses (Fig. 5A), we tested for signatures of functional segregation within supragranular layers, finding a strong bias for recurrent over cross-connections between L3a and L3b and a bias for connections from L2 to L3a over L3b. This descending connection is also more prevalent than the reverse ascending connection, from L3a to L2.

**Figure 5.**
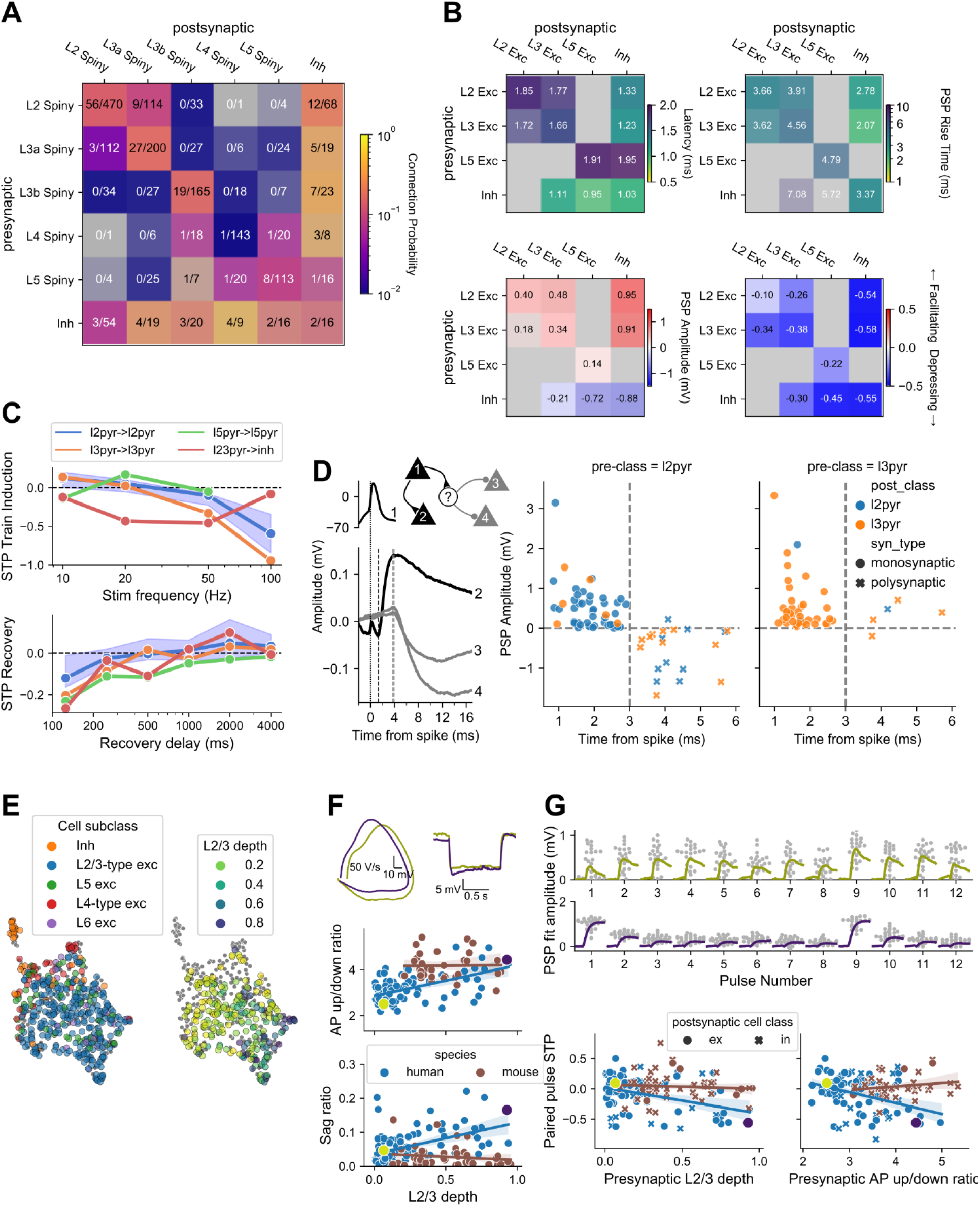
Human: **A.** Connection probability of human synapses. Inhibitory cells are identified by morphology as aspiny or sparsely spiny cells, grouped across layer. **B.** Kinetics, strength, and dynamics of human synapses. All matrices are organized by layer for excitatory cells, with inhibitory cells grouped across layer. Each element is colorized by the median (text in each element) according to the colormap with the saturation scaled to the standard error. Two or more pairs were required to fill in an element. Latency, Rise tau, and Resting state amplitude are quantified from fits of the average PSP response which have passed QC and visual inspection of the fit. **C**.Train induced STP (top) across frequencies for a subset of connection types. Each dot is the median of all synapses in the element, with shading for the 95% CI (bootstrapped) shown for a single example connection type. Lower plot shows recovery from STP at different delays. **D.** Example polysynaptic circuit from one experiment in which cell 1 forms a short latency (∼2ms) monosynaptic excitatory connection to cell 2 and delayed (∼4 ms) polysynaptic inhibitory connections to cells 3 and 4 (all cells confirmed morphologically spiny). Dashed lines indicate (from left to right) time of presynaptic spike and PSP onset. Polysynaptic connections from L2/3 pyramidal cells inferred by response latency > 3 ms are plotted with PSP amplitude. All polysynapses from L2 pyramidal cells (left) are inhibitory, while those from L3 pyramidal cells (right) are mixed. **E**. UMAP projection of intrinsic electrophysiology features from all human cells, colored by cell subclass (left), showing the distinctiveness of inhibitory and L4-type excitatory cells; and by depth (right), showing the structured variability of intrinsic properties with depth in L2/3. **F**. Within L2/3, variation in intrinsic properties is structured and strongly correlated with depth in human but not mouse. Example traces are shown for a superficial and deep human cell (colors as in D), alongside scatter plots of electrophysiology features vs. depth for both species (right). Top shows a phase plane representation of the first spike in a depolarizing step response, quantified as AP upstroke/downstroke ratio. Bottom shows the sag in response to hyperpolarization, quantified as sag ratio. Regression lines shown with bootstrapped 95% CI. **G**. STP of L2/3 excitatory synapses is structured by depth in human and not mouse. Top shows PSP responses to spike trains for example cells from E. The larger response to the first spike is quantified by paired pulse STP, plotted below in relation to presynaptic depth (left) and AP up/down ratio (right) (postsynaptic relationships shown in Fig. S7E).

Recurrent connectivity within human L4 (*p_max_*=0.01 [0.0, 0.04] 95% CI, 1/143) is significantly lower than observed in the mouse (*p_max_*=0.21 [0.15, 0.28], 45/464). This contrast could be related to age ^(39)^, species, or brain area. Further, other observed connections involving L4 pyramidal cells (e.g. I→E and E→I) suggest that low excitatory recurrence is not a technical limitation of our dataset, but rather reveals a unique property of the human L4 circuit.

### Human synaptic properties

The strength, kinetics, and STP properties of the human synapses show moderate differences across layers and large differences by cell class, largely resembling observations in mouse (Fig. 5B). Recurrent excitatory connections in human cortex (E→E) have longer latency than those with a pre- or postsynaptic inhibitory cell (E→E median 1.73 ms vs. I→E 1.03 ms, KS test p=2.2e^−5^; E→I 1.34 ms, p=8.3e^−4^), and PSP rise times are faster for E→I than E→E connections (2.48 ms vs. 4.11 ms, p=8.0e^−8^), but slower for I→E connections (6.33 ms; p=0.014, 3.2e^−6^ vs. E→E, E→I). We observe some differences between L2 and L3, including presynaptic L3 cells forming more depressing connections than L2 cells (STP ratios −0.41 vs.-0.15, p=7.0e^−3^). We also note that E→I synapses are uniformly depressing, consistent with the identification of those inhibitory cells as fast-spiking Pvalb cells.

In certain properties, we did observe contrasts between human and mouse synapses. The overall amplitudes of L2/3 E→E and E→I connections are significantly larger than the corresponding synapses in mouse, both excitatory resting state response (E→E 0.37 mV human vs. 0.10 mouse, p=0.039; E→I 0.93 mV vs. 0.27 mV, p=7.0e^−4^) and 90th percentile response (E→E 0.51 mV vs. 0.29 mV, p=0.040; E→I 1.17 mV vs 0.87 mV, p=0.27). We also observe dramatically faster recovery from STP in human than mouse excitatory synapses, with most fully recovered at 500 ms (Fig. 5C). This contrast has been previously noted in recurrent L2/3 excitatory synapses ^(25)^; our observations suggest that it holds for L5 E→E and L2/3 E→I synapses also.

#### Human polysynaptic events

Large-amplitude synaptic connections in human, but not mouse cortex, have been reported to trigger polysynaptic, complex events ^(56)^. Indeed, we also see polysynaptic events, primarily short-latency (∼3ms) inhibition (Fig 5D). Plotting latency versus PSP amplitude of human synapses (Fig 5D) reveals a clear boundary where responses with a latency of 3 ms or greater, evoked from a confirmed pyramidal cell, are almost exclusively inhibitory (median latency of IPSPs: 4.04 [3.77, 4.49] ms), compared to monosynaptic EPSPs which had a latency less than 3 ms (median EPSP latency: 1.7 [1.44, 2.13] ms). This potential disynaptic inhibition (dIPSPs) originates in L2 and projects to other L2 (n=10) or L3 (n=11) pyramidal cells, with just one ascending polysynaptic response originating in L3. This directionality is consistent with the directionality of monosynaptic excitation across layers 2 and 3 as well as supported by higher connection rates from L2 pyramidal cells to inhibitory cells. Median latency of recurrent L2 dIPSPs (4.08 [3.92, 4.56] ms) is similar to that of L2→L3 dIPSPS (3.84 [3.58, 4.24] ms), but recurrent L2 dIPSPs (−0.86 [−0.09, −1.03] mV) are almost three times larger than L2→L3 dIPSPs (−0.28 [−0.11, −0.44] mV).

#### Variation with depth in human layers 2 and 3

Another property of supragranular neurons observed in human, but not in mouse, is strong depth-driven variability of intrinsic electrophysiological properties ^(24, 57)^. Visualizing the electrophysiology feature space by a UMAP projection of 27 electrophysiology features (Methods), we found that L4-type cells (high input resistance) are situated around the perimeter, indicating distinct properties from L2/3-type cells (Fig 5E). For the L2/3-type cells, projecting a normalized layer depth coordinate (relative to the L2+L3 thickness) onto this space shows a mostly smooth gradient of electrophysiological properties with layer depth, also verified by direct examination of depth correlations with sag and AP upstroke/downstroke ratio (p=2.9e^−7^, 6.7e^−6^) (Fig 5F). As previously observed, this correlation is not found in mouse L2/3 cells (p>0.09 for both).

STP metrics revealed a similar linear variation with layer depth in synapses from L2/3 pyramidal cells across both E→E (n=85 human, n=19 mouse) and E→I (n=19 human, n=71 mouse) connections. The paired-pulse STP showed a strong linear relationship with depth in the human data (p=2.6e^−5^), varying from weak facilitation for the most superficial cells (0.03 ± 0.03, mean±SEM) to depression for the deepest (−0.37 ± 0.10). A strong correlation was also found with the action potential upstroke/downstroke ratio of the presynaptic cell only (p=2.2e^−5^ vs p>0.1 for postsynaptic), suggesting that links between spike shape and neurotransmitter release could help explain the variation of STP with depth (Fig 5G). No corresponding trends were found in the mouse data (p>0.1 for all regressions). Although lower sampling of L2/3 pyramidal cells may contribute, regression coefficients for STP against upstroke/downstroke ratio show a strong contrast (human −0.18 [−0.26, −0.10] CI; mouse 0.06 [−0.04, 0.16]), suggesting that there are real differences between these datasets in the factors contributing to STP variability, whether explained by species, brain area, or other factors.

### Modeling short term plasticity of mouse and human synapses

The STP metrics introduced above were chosen for their ease of interpretation, but were derived from a limited set of stimuli, and thus provide a simplistic description of the complex behaviors expressed by synapses. Ideally, we would like a description that can predict synapse behavior in response to any arbitrary stimulus. Rather than continue to engineer more descriptive statistics, we developed a generative model of stochastic vesicle release and STP with several adjustable parameters (see Methods) and asked which combinations of parameters were best able to explain the responses recorded for each synapse. In this way, we capture and describe more of the dynamic behavior of each synapse with a small number of parameters.

Model performance was evaluated by using the maximum likelihood parameter set for each synapse to simulate experimental data. This simulated data was then used to generate the same STP metrics that were previously collected from synaptic data. Both resting state PSP amplitude and 90th percentile amplitude are almost perfectly correlated between recorded and simulated data (Fig S12A,B), indicating that the model does exceptionally well at capturing synaptic strength. STP and variability (Fig S12C-F) are also strongly correlated, but with more scatter relative to strength metrics. Notably, STP measured from the second pulse in 50 Hz trains was only half as large as measured in synaptic responses, indicating that the model as parameterized was not able to fully capture STP on this timescale. Resting state variability is fit by the model through the binomial coefficient of variation, which is the product of the resting state release probability and the number of synaptic release sites. Release probability was more highly correlated with variance than number of release sites, suggesting that synapses may control variability primarily through their release probability.

Most discussions of short term depression in the cortical literature begin with the assumption that depression is caused by the depletion of vesicles from the readily-releasable pool. However, recent evidence suggests that calcium channel inactivation may be a more prominent mechanism in cortical depression ^(58)^. We ran the model on two separate parameter spaces--one that uses vesicle depletion, and another that uses a release-independent depression mechanism. In most cases, the model maximum likelihood value was found in the release-independent parameter space (Fig S11E, left). Release-dependent depression mechanisms should result in negative correlation between consecutive PSP amplitudes; however, we find little evidence for such negative correlations in our mouse data. Likewise, we find little relationship between paired correlation values and the model preference for release-dependent mechanisms (Fig S11E, right). These results are consistent with the proposal that cortical synapses in mouse employ release-independent depression mechanisms, and that vesicle depletion plays a relatively minor role in depression. In comparison, our data from human synapses does have a modest preference for negative correlation between paired event amplitudes.

### Organization in mouse and human synaptic dynamics

With a large dataset describing synapse properties it becomes possible to ask what patterns emerge from the data. What synaptic features correlate with one another, and do synapses naturally split into clusters based on these features, or do they form a continuum? What aspects of the synaptic feature-space are driven by presynaptic versus postsynaptic cell type? Prior studies have found that excitatory synapses onto Pvalb and Sst cells have distinctly different dynamics, suggesting a general rule that excitatory dynamics depend primarily on the postsynaptic cell type ^(59, 60)^. In contrast, inhibitory dynamics have been found to depend mainly on the presynaptic type, particularly when comparing Pvalb to Sst ^(61, 62)^. Although our data often follow these rules, we also find exceptions and suggest some refinements.

We used sparse PCA followed by UMAP dimensionality reduction to summarize the output of the stochastic release model described above for 1140 synapses (980 mouse, 160 human) (Fig. 6). This analysis groups synapses based on the similarity of their model results; therefore, it has access to any synaptic strength and dynamical properties that the model could capture, but does not have access to any information about kinetics, cell subclass, or other cell properties.

**Figure 6:**
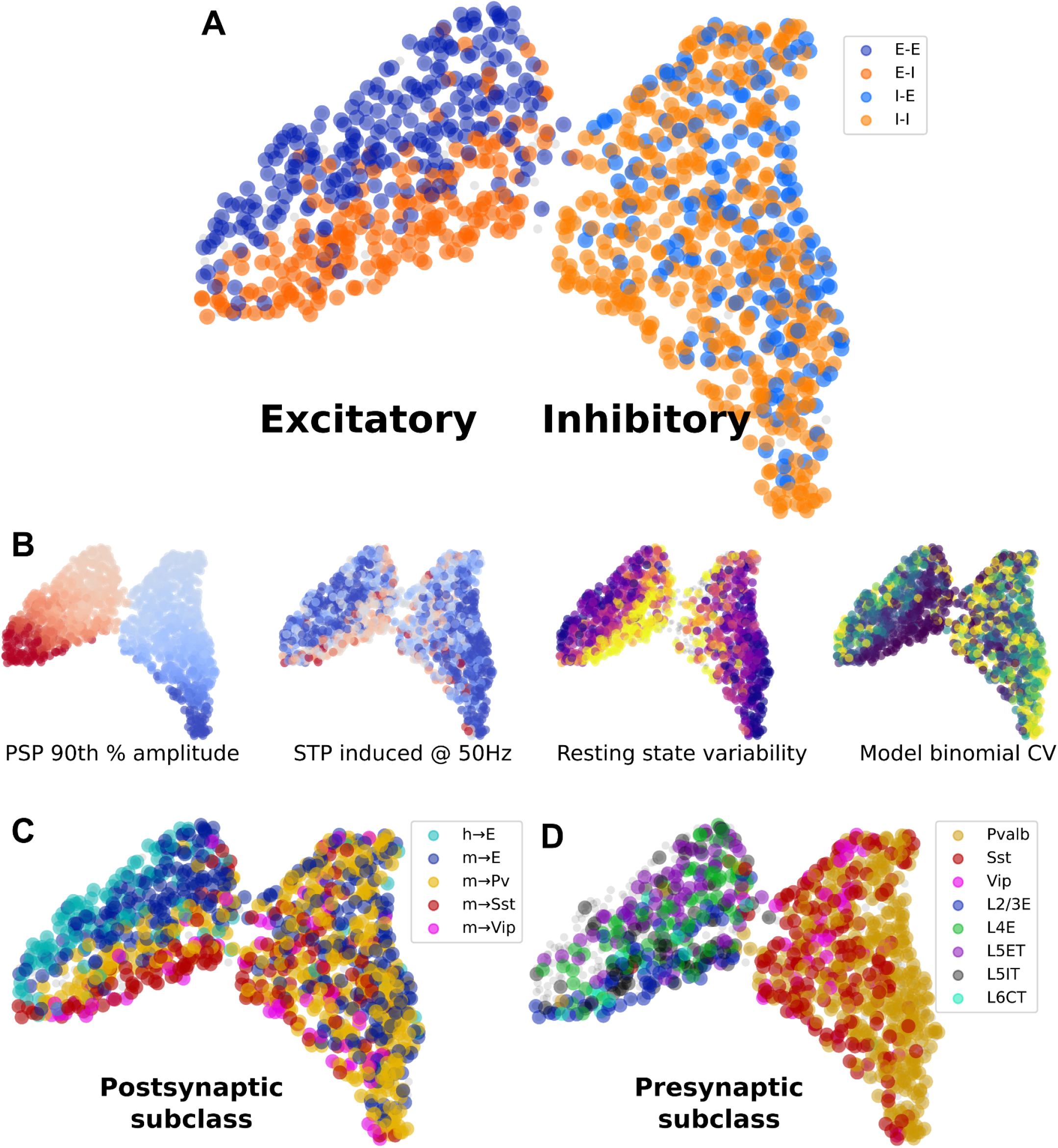
Relationships among synapse properties and cell types revealed by dimensionality reduction. **A.** All synapses colored by postsynaptic E/I cell class. The UMAP output generates two clusters: excitatory (left) and inhibitory (right). **B.** Four synapse properties represented in reduced space, showing 90th percentile PSP amplitude (red=excitatory, blue=inhibitory); STP induced by 50 Hz trains (red=facilitating, blue=depressing), resting state aCV during 50 Hz trains (purple=low variability, yellow=high variability), and the binomial CV derived from model parameters (release probability * number of release sites; purple=high CV, yellow=low CV). **C.** Human and mouse synapses colored by postsynaptic subclass. **D.** Mouse synapses colored by presynaptic subclass.

Synapses in this analysis form clear excitatory and inhibitory clusters (Fig. 6A), with a continuum of synaptic properties within each cluster. Perhaps the most prominent feature of this organization is that excitatory synapses are strongly differentiated by the postsynaptic E/I class, whereas inhibitory synapse properties are mostly independent of postsynaptic cell class.

What synaptic properties determine this organization by class? The most clear property that correlates with the UMAP dimensions is synaptic strength, which forms a well defined gradient with the long axis of each cluster (Fig. 6B, left). However, this gradient is mostly parallel to the boundary separating cell classes, indicating that cell classes are not strongly differentiated by strength. Orthogonal to the strength axis, we find that response *variability* differentiates synapses and strongly separates postsynaptic classes in the excitatory cluster. This axis also correlates with STP, where the most facilitating synapses appear at the high-variability end of the axis.

The excitatory cluster is further stratified by postsynaptic cell subclass, with the most distinct separations seen between E→Sst, E→Pvalb, mouse E→E, and human E→E (Fig. 6C). In this context we see that Sst and Vip cells receive the highest-variance (and also most facilitating) synapses, whereas human E→E synapses are distinctly more reliable, compared to mouse. In contrast to these results, excitatory synapses seemed only weakly differentiated by the presynaptic subclass. These results confirm the rule that excitatory synaptic dynamics are largely determined by the postsynaptic cell.

Most inhibitory synapses in this analysis occupy a central part of the inhibitory cluster that is relatively independent of either pre- or postsynaptic subclass (Fig. 6C,D). This region is characterized by moderate variability and a range of STP from weakly depressing to facilitating. One major exception is that a subset of Pvalb synapses occupy the far edge of the inhibitory cluster almost exclusively (Fig. 6D, right), where variability is low and STP is strongly depressing. However, Sst synapses are somewhat differentiated by their postsynaptic cell type, especially when connecting to Vip cells. This suggests that unlike excitatory dynamics, inhibitory dynamics do not follow a simple rule related to either pre- or postsynaptic subclass.

## Discussion

The mammalian cortex is believed to act as the computational substrate for our highest cognitive abilities, particularly the ability to model the world around us and predict the effects of our actions. It is also of particular interest because many aspects of its structure are repeated across brain regions and conserved across species, suggesting the existence of a general-purpose approach to computation in the cortex. There is a long history of electrophysiological and anatomical experiments exploring the local connectivity of the cortical microcircuit. Microcircuit representations have evolved from experiments in different species, regions, ages, etc that focus on one or a few circuit elements. These efforts offer an excellent depth of insight to isolated regions of the circuit but lack a complete and unified view of the circuit ^(14)^. Furthermore the difficulty of accessing these historical data discourages reuse and reanalysis. We saw an opportunity to expand upon this history and conduct a broader survey than has been attempted in the past, and extend the value of this resource to the community by making our analyses, tools, and data open to the public.

By probing over 20,000 possible connections across 28 mouse lines, we have explored a large fraction of the subclass-specific, intralaminar connectivity in the mouse visual cortex. At the same time, we share the first systematic survey of intralaminar connectivity in the human cortex. Past surveys near this scale have focused on connectivity and strength of synapses; a major advance provided by our study is the depth of characterization and analysis for each synapse, in the context of transgenically identified cell subclasses and species.

### A proposed standardized model of connectivity

The likelihood that two neurons are locally connected depends on multiple factors such as cell type, cortical region, species, and animal age. A longstanding goal has been to determine the governing principles of local circuit architecture^(4, 6, 50, 63–66)^. It is difficult to make direct comparisons among these studies because the observed rate of connectivity depends on several experimental details that are often inadequately reported or controlled for. For example, the probability of connection between two cells depends strongly on the intersomatic distance ^(27, 67)^; thus an experiment that samples shorter intersomatic distances is likely to report a higher rate of connectivity, even for the same ground-truth circuit. Likewise, false negatives may occur when measuring connectivity as a result of severed connections or poor signal detection power.

Ideally, we would like a way to describe connectivity that accounts for these effects (and potentially others), allowing more direct comparison between experiments regardless of their methodological differences. In order to facilitate such comparison between connection subclasses in our own data, we developed a procedure for modeling connection probability as it relates to intersomatic distance, axon truncation, cell depth, and signal detection power. With this model, we can estimate unbiased connection probabilities with confidence intervals that should be relatively robust to experimental bias. In principle, this approach is flexible enough to be replicated elsewhere in the field and its adoption would substantially improve our ability to compare and reproduce results across studies.

### Conserved and canonical elements in the mouse intralaminar circuit

As many prior studies have investigated the cortical circuit, a picture has emerged describing the relationships between the excitatory and inhibitory subclasses and their functional relevance. The details of this picture vary somewhat between descriptions, but a few key elements appear consistently, especially in the systems and theoretical neuroscience literature (Fig 7A). Pvalb interneurons strongly inhibit nearby pyramidal cells and other Pvalb cells, Sst interneurons broadly inhibit nearby cells but avoid other Sst cells, and Vip cells selectively inhibit Sst cells and receive feedback excitation, forming a disinhibitory circuit ^(35, 36)^. We have confirmed that each of these motifs is prominent across layers in the intralaminar cortical circuit.

**Figure 7:**
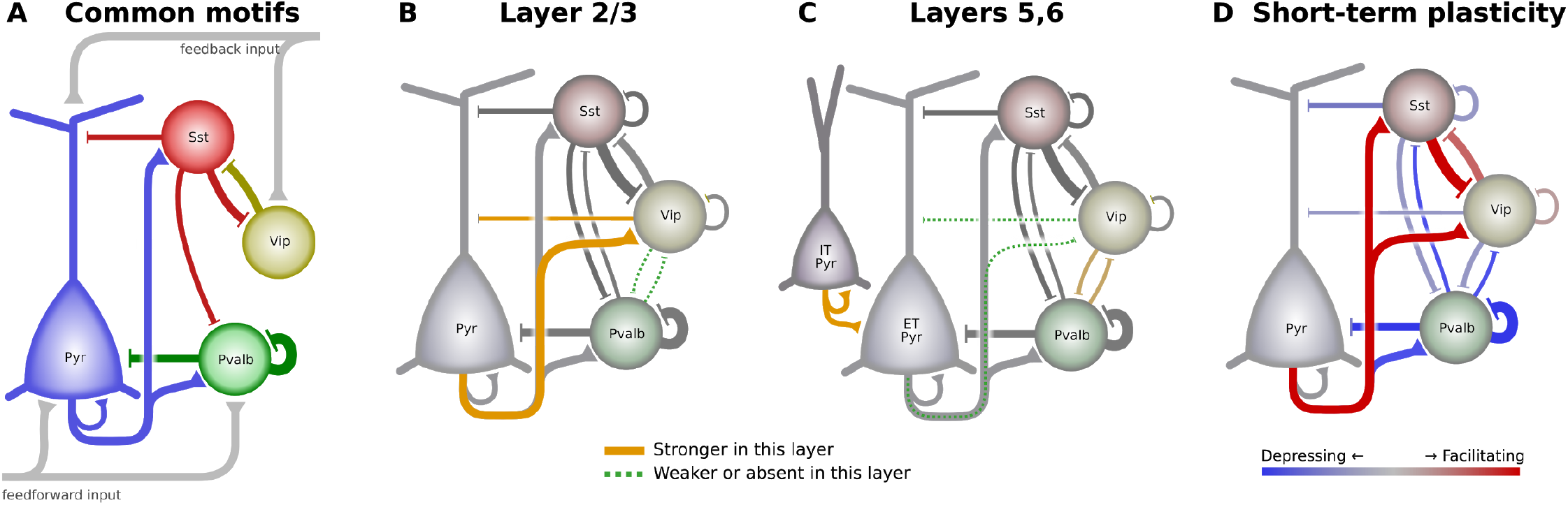
The cortical intralayer circuit differs across layer and with activity. **A)** Some commonly described elements of the intralaminar cortical circuit. Pvalb cells strongly inhibit pyramidal and other Pvalb cells, Sst cells provide broad inhibition, and Vip cells inhibit Sst cells to form a disinhibitory feedback pathway. **B)** Layer 2/3 circuit diagram showing connections between major subclasses. The width of connecting lines roughly represents connection probability and PSP amplitude. Connections that are prominent in mouse L2/3 compared to deeper layers are highlighted in orange, whereas green dashed lines indicate connections that are weak or absent in L2/3. **C)** Circuit diagram of connections found in mouse layers 5 and 6. For simplicity, connections between IT pyramidal and inhibitory cells are omitted. **D)** Two complementary circuits that activate at different times. Red connections are facilitating and will be stronger during sustained activity. Blue connections are depressing and are strongest during quiescent periods.

We also find circuit elements that are equally prominent in our data, but are more sparsely acknowledged in the literature. Many recent studies have focused on the importance of the Vip→Sst disinhibitory circuit. In the opposite direction, however, the connection from Sst to Vip has one of the highest connection probabilities, largest IPSP amplitudes, and strongest facilitation in our dataset. Although Sst→Vip connections have been described previously ^(7)^, they are often overlooked in consideration of the disinhibitory circuit. The unique synaptic features we observed suggest an important functional ramification on the opposing Vip→Sst disinhibitory pathway. For example, local excitation could drive disinhibition either through the established Vip→Sst pathway or through the reverse Sst→Vip pathway. Furthermore, mutual inhibition between Vip and Sst populations could result in bistability in which either subclass may exclusively drive inhibition of pyramidal cells, but not both simultaneously.

Sst and Vip cells are often described as lacking recurrent connections ^(11, 39, 54, 68)^ despite some evidence to the contrary ^(44, 69)^. We have confirmed that Sst and Vip do have the expected biases in their connectivity across all layers (Sst cells tend to avoid contacting other Sst cells and Vip cells prefer to contact Sst cells). However, we also find sparse, recurrent connections within both interneuron populations about equal to the recurrent connectivity in excitatory populations. Furthermore, we find that the strength of connections from Sst and Vip do not follow the same preferences, having roughly equal strength when connecting to preferred versus non-preferred subclasses. Given their trans-laminar axon projection patterns, recurrent connections within these subclasses may be found more commonly across layer boundaries ^(13)^.

### Laminar variations on the cortical circuit

Previous studies that sampled both L2/3 and L5 noted many similarities between the two layers ^(4, 6)^. Although we have confirmed a consistent set of connectivity rules describing the intralaminar circuit, we also find variations on these rules that could support different modes of cortical function. Differences in intralaminar circuitry may contribute to laminar differences in receptive field properties ^(70)^ or visually mediated behaviors ^(36)^.

Layer 2/3 has strong interconnections between pyramidal and Vip cells (Fig 7B). In deep layers, these connections are either absent or greatly reduced, and the relative sparsity of Vip cells in deep layers should further enhance this difference ^(71, 72)^. Direct Vip inhibition of local pyramidal cells further complicates the prevailing view of the Vip→Sst disinhibitory pathway by, for example, allowing the possibility of feedback inhibition from higher cortical regions. Likewise, local excitatory inputs to Vip cells could be a source of feedforward disinhibition in layer 2/3. In contrast to L2/3, recurrent connections between the Pvalb and Vip subclasses were found most frequently in deep layers (Fig 7C). Ultimately, more targeted experiments and modeling will be needed to explore the functional relevance of these laminar variations in the cortical circuit.

Layer 5 excitatory subclasses differ in their visual responses and long range projections, suggesting different functional roles in the circuit ^(73, 74)^. In this context we see several interesting differences in L5 ET and IT neurons. ET pyramidal cells are generally more highly connected, receiving more local excitation and inhibition than IT cells. We confirmed a much higher rate of recurrent connections among L5 ET cells compared to IT as well as the observation that connections between these two subclasses are unidirectional from IT→ET (Fig 7C) ^(37)^. Layer 5 ET cells also receive more frequent inhibition from Sst cells, as previously observed in frontal cortex of rat ^(38)^.

Could laminar differences in connectivity indicate cell type divisions within subclasses? Two recent studies investigated the correspondence between morphological, electrophysiological, and transcriptomic (MET) features of inhibitory neurons in primary visual cortex and motor cortex ^(13, 75)^. In visual cortex, different MET types had distinct patterns of local axonal innervation and dendritic morphologies ^(13)^ suggesting that their connectivity will be different. MET types also exhibited layer localization and thus some of the differences in connectivity we observe as a function of layer may reflect differences in connectivity between different MET types.

### Dynamic flexibility in the cortical circuit

Most studies in cortical synaptic physiology describe the circuit in its quiescent state. Ongoing activity *in vivo*, however, dynamically rewires the network by strengthening or weakening synapses. Cortical up- and down-states in particular have the capacity to synchronously facilitate or depress large portions of the network. We find a diversity of dynamic properties even among specific cell subclasses; thus to some extent this dynamic network reshaping occurs at the level of individual synapses. However, we also find that many subclass elements of the cortical circuit show a clear preference for either facilitation or depression (Fig 7D). Overall, most synapses in our dataset were found to exhibit synaptic depression. Pvalb cells in particular receive and project almost exclusively depressing synapses. In contrast, Vip and Sst connections express a mixture of depression and facilitation. Excitatory inputs to these subclasses are often strongly facilitating, as are the interconnections between Vip and Sst. These patterns suggest the ability to dynamically switch between two network modalities where intralaminar activity is either dominated by facilitating interactions between pyramidal, Sst, and Vip cells during sustained activity, or dominated by interactions between pyramidal and Pvalb cells at the onset of activity.

### Excitatory cells receive weak local excitation compared to inhibitory cells

Our previous study ^(51)^ observed low recurrent excitatory connectivity rates in all layers. We now find that recurrent excitatory synapses have a set of features that distinguish them from excitatory inputs to inhibitory cells and appear to limit their contribution to excitability. In addition to being sparse, they are relatively weak, they get weaker with activity, and they have slower PSP rise times. Slow PSP rise times would limit excitability by raising action potential threshold (Azouz and Gray, 2000). Recurrent excitatory connections also have relatively long latencies. A well-described feature of cortical networks is that inhibition lags behind excitation. This correlation has wide-ranging functional consequences, including creating windows of integration (Pouille and Scanziani, 2001; Pouille et al., 2009). It is interesting to consider the differences between mouse visual cortex and human temporal cortex recurrent excitatory rates in this context. L4 in mouse has the highest recurrent connectivity rate of all layers. L4 also receives potent feedforward inhibition in the visual cortex ^(76)^ that may coincide with the long latency recurrent excitation to shape the integration window. In human temporal cortex, recurrent excitation is practically absent in L4,and thus is unlikely to shape the integration window.

### Unidirectional disynaptic inhibition in human

We observe disynaptic inhibition in human cortex between confirmed spiny pyramidal cells that is unidirectional, originating in L2 and targeting other L2 or L3 pyramidal cells. We did not see disynaptic inhibition in our mouse recordings, which may be due to the stronger excitation we, and others ^(56)^, observe in human synapses, particularly onto inhibitory cells. Disynaptic inhibition is often mediated by an interposed Sst cell ^(77, 78)^ as they have a low spiking threshold and receive facilitating inputs from excitatory cells. However, the latency of Sst-mediated disynaptic inhibition is often long (> 100 ms) whereas we saw disynaptic IPSPs with a much shorter latency (3 - 6 ms). This suggests the disynaptic inhibition is driven by an intermediate fast-spiking Pvalb cell which has been observed in human and can be recruited by very large excitatory events ^(79)^. The unidirectional nature of this disynaptic inhibition from more superficial to deeper cortex further suggests a preferential routing of information by Pvalb cells in human cortex ^(79)^.

### Synapse types differ in variability

We have taken two complementary approaches to describe the dynamic behavior of mouse and human synapses. In the first, we take advantage of repeated stimuli of varying temporal structure in order to measure the effects of short-term plasticity and quantal variance. These metrics are generally easy to interpret but have several drawbacks: they are sensitive to noise, they require successfully repeated stimuli that are not available for all synapses, and they are difficult to use in a biophysical modeling context. In the second approach we address these issues with a new generative model describing quantal release and STP. With this model we are able to use all response data available for each synapse and yield a consistent set of parameters with a clear path to model implementation. Our approach is similar to other recently developed models ^(80, 81)^ in that it does not depend on any particular stimulus structure (aside from having a diversity of interspike intervals), and thus frees the experimenter to design stimuli by other criteria and gracefully handles quality issues such as spike failures and early experiment termination.

The model output for each synapse is a multidimensional map of the model likelihood measured across a large parameter space. Each map contains a signature that is unique to the synapse, but shares some features with similar synapses. By reducing the dimensionality of this model output, we were able to ask whether there exists a natural organization among the different types of synapses across both species in our dataset. A few governing principles emerge from this analysis. We find that cortical synapse types can be organized into a two-dimensional feature space, with synaptic strength and variability forming two orthogonal gradients. For the most part, synapses occupy a continuum across these two features. Short-term plasticity features also vary in parallel with variability, but in a discontinuous manner compared to the smoother gradient of variability. In the reduced-dimensionality map, we find that excitatory synapse variability is strongly differentiated across postsynaptic cell type. Sst synapse dynamics also differ by postsynaptic subclass, Vip synapse dynamics are largely independent of the postsynaptic subclass, and Pvalb synapses on average have much less variability than other inhibitory types, regardless of the postsynaptic subclass. The rules derived from these relationships elaborate on simpler rules described previously ^(61, 62, 82)^.

Synaptic variability has, in the past, been regarded as an undesirable consequence of signaling via metabolically expensive exocytosis. More recently, advances in machine learning that rely on stochasticity have supported the possibility that synaptic variability may offer computational benefits, such as a mechanism for regularization during learning^(21)^. If that is the case, it is further plausible that variability may be modulated by cell type, and that these relationships are crucial features of cortical function. Indeed, our measurements of synaptic variability were found to strongly differentiate cell type in a pattern that is largely, but not entirely, aligned with STP metrics, suggesting the possibility of cell type-specific tuning of variability. More broadly, analysis of our stochastic release model indicates that synaptic strength and variability form the two most significant parameters describing synapse behavior. It remains to be explored whether variability itself is an objective parameter directly tuned by the cortex, or if this is a consequence of the relationship between variability and short term plasticity.

## Limitations

The methods we use require brain tissue to be dissected and sliced to provide easier visual and physical access to the neurons to be patched. In this process, some neurons are damaged, some connections are severed, and much of the *in vivo* environment is replaced with controlled conditions. At the same time, the synaptic effects we are interested in can be measured only indirectly via their impact on the cell soma. Our ability to resolve these small signals is affected by dendritic filtering, background noise, and the inherent variability of the synapse. Despite these limitations, paired patch-clamp recording remains the gold standard in synaptic physiology for its excellent electrical resolution, temporal precision, and control over presynaptic spiking. Past studies have modeled the effect of severed connections ^(27)^ and sensitivity ^(51)^ on the rate of false negatives in connectivity measurements. We believe this is the first study to quantify these sources of error simultaneously and estimate adjustments, which are substantial, especially for mouse inhibitory connectivity. In addition to providing better estimates of connectivity, reducing experimental bias makes it possible to compare our results between studies with more confidence. For example, recurrent excitatory connections were found more frequently in human, but our analysis of detection power suggests that this difference is due to enhanced signal to noise ratio (arising from a lower noise floor and stronger and more reliable PSPs) in human cortical tissue.

A confounding factor associated with human neurosurgical tissue is the potential for disease pathology (epilepsy or tumor in this case) to affect the physiological measurements we are recording such that we may not be capturing the healthy human brain. While this is important to keep in mind, these tissues are currently our only access to living human cortex. Additionally, a recent study ^(24)^ on intrinsic cell physiological features in human found strikingly consistent results across nearly 100 different human tissue donors who varied in age, gender, cortical region, and pathology. This suggests that despite disease state, we can gain useful insight into human cortical function through the use of neurosurgical specimens.

For this study, we focused our efforts on intralaminar connectivity; instances of interlayer connections are present in the dataset but relatively sparsely sampled. For nearby layers, this could be addressed with further experiments. However connections across distant layers are more difficult to interrogate with this method because long axo-dendritic pathways are more likely to be severed ^(32)^, especially in human cortical tissue that can be several millimeters from pia to white matter. A detailed study of such medium-range connections may require a different methodological approach ^(83)^.

The transgenic mouse lines used in this study enabled the investigation of connectivity and synaptic physiology between subclasses that are otherwise difficult to target in slice recordings. However, they also merge more refined cell types ^(13, 84)^ and miss others. The most notable omission is transgenic mice that identify serotonin positive, Vip negative interneurons which make up about 20% of the interneuron population ^(8)^. Merging cell types is a common issue when a single modality is used to assign a cell identity ^(85)^. To overcome this limitation, we provide a multimodal dataset (physiological and morphological), and these complementary modalities may be used to refine or verify the cell identity. End-users may utilize the various modalities in our dataset in ways that are appropriate for their research question.

### Future work

In the near term, we are excited about the incorporation of our data into biophysically realistic network models to address questions related to cell-type and synaptic dynamics. With the recent advancements in the segmentation and annotation of large volumes from electron-microscopy ^(86)^, the opportunity to reconcile connectivity measurements from functional studies like ours with anatomical measurements is on the horizon. The combination of patchSeq ^(13, 41)^ and multipatch experiments creates the opportunity to gain insights into local connectivity among transcriptomic cell types. A straightforward path is the continuation of the pipeline to explore other brain regions’ microcircuitry to reveal the extent to which circuits are either conserved or specialized. A deeper understanding of brain function will require data on the relationships between cell types, learning rules, and modulatory pathways.

## Acknowledgements

The authors wish to acknowledge the founder of the Allen Institute, Paul G. Allen, for his vision, encouragement and support.

## Materials and methods

Methods were similar to Seeman, Campagnola et. al, 2018. Information on the Synaptic Physiology pipeline and the dataset are accessible from our website (Synaptic Physiology Coarse Matrix Dataset). Full Standard Operating Procedures can be found: https://www.protocols.io/workspaces/allen-institute-for-brain-science/publications?categories=multipatch

### Animals and tissue preparation

All animal procedures were approved by the Institutional Animal Care and Use Committee at the Allen Institute for Brain Science (Seattle, WA), which operates per National Institutes of Health guidelines. Triple (T.L.D., unpublished) and quadruple ^(26)^ mouse lines generated using double transgenic mouse lines, were used to target up to two unique cell subclasses in a single animal (see Table S1, http://portal.brain-map.org/explore/toolkit/mice). Each subclass was selectively labeled by fluorescent reporters (tdTomato or EGFP) driven by Cre or FlpO. Layer-specific excitatory cells were targeted using unique transgenic drivers: Nr5a1 and Rorb for layer 4, Sim1, and Tlx3 for layer 5 ET and IT, respectively, and Ntsr1 for layer 6 CT. It was generally not possible to generate crosses of two excitatory drivers, however in L5 we were able to target ET cells via a retroorbital injection of mscRE4-FlpO AAV PHPe.B ^(87)^ into Tlx3-Cre transgenic mice in order to probe interconnections of L5 ET and IT cells. Transgenic lines were not used to target layer 2/3 excitatory cells but were later confirmed through the presence of dendritic spines via post-hoc morphological analysis (see [Morphology and Position](https://docs.google.com/document/d/1d1YRsw7Yu30fLHBYdd8jZmsdAVYo4f9n92odLl5LOes/edit#heading=h.a1htosvo9dl7)). Inhibitory cell subclasses, Sst, Pvalb, and Vip, were targeted in all layers.

Female and male adult mice (mean age 46.0 ± 4.6; SD) were anesthetized with 5% isoflurane and transcardially perfused with ice-cold oxygenated slicing aCSF I. All aCSF recipes are in Table S2.

Acute parasagittal slices (350 µm) were produced with a Compresstome (Precisionary Instruments) or VT1200S Vibratome (Leica Biosystems) in ice-cold aCSF I solution. The slicing angle was set to 17° relative to the sagittal plane to preserve pyramidal cells’ apical dendrites. Slices were then recovered for 10 min in a holding chamber containing oxygenated aCSF I maintained at 34°C. After recovery, slices were kept in room temperature oxygenated aCSF IV (Table S2).

Human neocortical tissue from Temporal, Frontal, and Parietal lobes was obtained from adult patients undergoing neurosurgery for the treatment of epilepsy (52 samples) or tumor (20 samples; Fig 1A). Tissue obtained from surgery was distal to the core pathological tissue and was deemed not to be of diagnostic value. Surgical specimens were placed in a sterile container filled with pre-chilled (2-4°C), carbogenated aCSF VII containing decreased sodium replaced with NMDG to reduce oxidative damage (Table S2), and delivered from the surgical site to the laboratory within 10-40 min.

In the laboratory, specimens were trimmed to isolate regions of interest and mounted to preserve intact cortical columns (pial surface to white matter) before being sliced in aCSF VII using a Compresstome or Vibratome. Slices were then transferred to oxygenated aCSF VII (34°C) for 10 min, then moved and kept in aCSF VIII at room temperature (Table S2) for a minimum of one hour prior to recording.

### Electrophysiological recordings

Slices were placed in custom recording chambers perfused (2-4 mL/min) with aCSF IX which contained one of two external calcium concentrations ([Ca^++^]_e_) 1.3 mM or 2.0 mM (Table S2). aCSF IX in the recording chamber was measured at 31-33°C, pH 7.2-7.3, and 30-50% oxygen saturation. In our previous study ^(51)^, we conducted experiments in mouse with a [Ca^++^]_e_ of 2 mM to be consistent with previous connectivity studies ^(6, 67, 88, 89)^. However, external calcium concentration *in vivo* has been measured to be closer to 1 mM ^(90)^ and more closely reproduces *in vivo*-like short term plasticity *in vitro* ^(91)^. Thus, we reduced [Ca^++^]_e_ to 1.3 mM to measure synaptic properties closer to physiological conditions. However, when we compared connection probability, strength, and short-term plasticity for connection elements in which we had data at both [Ca^++^]_e_ (21 elements for connectivity and 10 elements for synaptic properties out of 89 targeted intralaminar elements), we found connectivity and synapse characteristics were consistent between the two concentrations (Fig S8). Thus for results reported in this study, data were pooled across conditions. Experiments on human tissue were conducted with 1.3 mM [Ca^++^]_e_ only.

Recording pipettes (Sutter Instruments) were pulled using a DMZ Zeitz-Puller (Zeitz) to a tip resistance of 3-8 MΩ and filled with internal solution containing (in mM): 130 K-gluconate, 10 HEPES, 0 (human) or 0.3 (mouse) ethylene glycol-bis(β-aminoethyl ether)-N,N,N’,N’-tetraacetic acid (EGTA), 3 KCl, 0.23 Na2GTP, 6.35 Na2Phosphocreatine, 3.4 Mg-ATP, 13.4 Biocytin, and either 50 µM Cascade Blue dye (excited at 490 nm), or 50 µM Alexa-488 (excited at 565 nm).

Internal solution was measured with osmolarity between 280 and 295 mOsm with pH between 7.2 and 7.3. All electrophysiological values are reported without junction potential correction. We removed EGTA from our internal solution for human recordings to be consistent with previous human electrophysiological studies ^(56, 92)^. A small subset of human recordings were conducted with 0.3 mM EGTA. A comparison of connection elements in which we had both EGTA conditions showed consistent connectivity and synaptic properties and thus the data was pooled (Fig S9).

Eight recording headstages were mounted in a semi-circular arrangement around the recording chamber. The pipette holders were fitted with custom shields to reduce crosstalk artifacts. Each headstage was independently controlled using modified triple-axis motors (Scientifica; PatchStar). Recorded signals were amplified (Multiclamp 700B, Molecular Devices) and digitized (50-200 kHz) using ITC 1600 DAQs (Heka). Pipette pressure was controlled using electro-pneumatic control valves (Proportion-Air; PA2193) or, though manual, mouth applied pressure, available for one pipette at a time. Slices were visualized using oblique (Olympus; WI-OBCD) infrared illumination using 40x or 4x objectives on a custom motorized stage (Scientifica) using a digital sCMOS camera (Hamamatsu; Flash 4.0 V2). Acq4 software (acq4.org; ^(93)^) was used for pipette positioning, imaging, and subsequent image analysis.

Eight neurons (excitatory or inhibitory) were targeted based on cortical layer, somatic appearance, and depth from the slice’s surface in experiments from human and mouse tissue. Neurons in transgenic mice were also targeted based on fluorescent reporter expression. Cells were targeted with a depth of at least 40 µm (Fig S1B) from the surface of the slice with automated pipette control assistance. In order to minimize tissue distortion and damage, pipettes moved through the tissue on a trajectory that was collinear with the long axis of the pipette with minimal positive pressure (10 - 40 mBar).

Whole-cell patch-clamp electrophysiological recordings were performed on neurons that formed a stable seal and had a successful break-in. At least two neurons were measured at the same time per recording, with the mean number of simultaneous recordings being 4 for both mouse and human (see Fig 1A for distributions). Recordings were performed with a holding potential set to either −70 mV (to measure excitatory inputs) or −55 mV (to measure inhibitory inputs) and were maintained within 2 mV using automated bias current injection. Data acquisition was collected using Multi-channel Igor Electrophysiology Suite (MIES; https://github.com/AllenInstitute/MIES), custom software written in Igor Pro (WaveMetrics). A 15-18 second intersweep interval to allow the synapse to recover was used. During a sweep, evoked spikes were distributed in time across recordings such that they were separated by at least 150 ms.

To examine short-term plasticity (STP), cells were stimulated in current and voltage-clamp to drive trains of 12 action potentials (Fig 1B) at different fixed frequencies of 10, 20, 50, 100, and 200 Hz with a delay period between the 8th and 9th pulses ^(94)^. The delay period lasted 250 ms for all frequencies with additional delay periods (125, 500, 1000, 2000, 4000 ms) for 50 Hz stimulation. Protocols were repeated five times for each stimulation frequency and delay interval. We also delivered a “mixed frequency” stimulus which was composed of 8 action potentials at 30Hz immediately followed by 30 action potentials, whose intervals were a random resequencing of 29 exponentially increasing intervals between 5 and 100 ms. The intervals were fixed across sweeps and experiments. While in current-clamp, an additional set of stimuli was used to characterize intrinsic properties of each cell (Fig 1C). To estimate input resistance of the cell, a 1-second-long hyperpolarizing square pulse was delivered at an initial amplitude of −20 pA while keeping the neuron at −70 mV. The voltage response to each current step was measured online and successive current steps were titrated to target response voltages of −68, −72, −75, −80, and −85 mV so as to reliably activate I_h_ when present. To measure spiking properties, a long (500 ms) depolarizing square pulse stimulus was delivered that started at rheobase and increased 25 pA for 6 intervals. Lastly, we delivered a 15-second sinusoidal chirp that increased in frequency from 0.2 to 40 Hz and evoked a response magnitude that measured ∼10 mV from peak to trough.

### PatchSeq recordings and processing

PatchSeq recordings were performed in a subset of mouse experiments. To avoid sample contamination, surfaces, equipment, and materials were cleaned using DNA away (Thermo Scientific), RNAse Zap (Sigma-Aldrich), and nuclease-free water (in that order). aCSF V was made daily and filtered before use. Materials used to make and store aCSF V were cleaned thoroughly before use. Recording pipettes were filled with ∼1.75 µL of RNAse Inhibitor containing internal solution: 110 mM K-Gluconate, 4 mM KCl, 10 mM HEPES, 1 mM adenosine 5’-triphosphate magnesium salt, 0.3 mM guanosine 5’-triphosphate sodium salt hydrate, 10 mM sodium phosphocreatine, 0.2 mM ethylene glycol-bis (2-aminoehtylether)-N,N,N’,N’-tetraacetic acid, 20 µg/mL glycogen, 0.5 U/µL RNase Inhibitor, 0.5 % biocytin, and either 50 µM Cascade Blue dye (excited at 490 nm), or 50 µM Alexa-488 (excited at 565 nm).

In patchSeq experiments, a subset of stimuli were collected to limit progressive cell swelling associated with the addition of RNAse to the internal solution ^(41)^.

Methods for nuclei extraction and processing are similar to previous patchSeq studies ^(13, 41)^. At the end of the experiment, pipettes were adjusted to the soma center or placed near the nucleus, if visible. A small amount of negative pressure (∼0.5 psi) was applied to all pipettes simultaneously for cytosol extraction. Extraction time varied for each cell; pipettes were slowly (∼0.3 µm/s) retracted in the x and z-axis once the soma had visibly shrunk and/or the nucleus was visible at the tip of the pipette. Once pipettes were out of the slice, cytosol and/or nucleus content in each pipette were expelled into individual PCR tubes containing 11.5 µl of lysis buffer (Takara, 634984) and stored in −80 °C.

We used the SMART-Seq v4 Ultra Low Input RNA Kit for Sequencing (Takara, 634894) per the manufacturer’s instructions to reverse transcribe RNA and amplify full-length cDNA. To obtain detailed methods, see http://celltypes.brain-map.org, “Transcriptomics Overview Technical White Paper”. We identified transcriptomic types by mapping our Patch-seq transcriptomes data in the same methods mentioned in previous studies ^(13, 95)^.

### Histology and imaging

After electrophysiological recordings, slices were fixed in solution containing 4% paraformaldehyde and 2.5% glutaraldehyde for at least 40 hours at 4°C. Slices were then transferred and washed in phosphate buffer saline (PBS) solution for 1-7 days before staining.

A 3,3’-diaminobenzidine (DAB) peroxidase substrate kit (Vector Laboratories) generated a brown reaction product in biocytin-filled neurons. Slices were stained with 5 µM

4’,6-diamidino-2-phenylindole (DAPI) in PBS for 15 min at room temperature and then triple-washed in PBS (10 min for each wash). Slices were then transferred to 1% hydrogen peroxide (H2O2) in PBS for 30 min and triple washed in PBS. Afterward, slices were mounted onto gelatin-coated slides and cover-slipped with Aqua-Poly/Mount (Polysciences). Slides were dried for approximately 2 days before imaging.

Mounted slides were imaged on an AxioImager Z2 microscope (Zeiss) equipped with an Axiocam 506 monochrome camera. Tiled mosaic images of whole slices were captured with a 20x objective lens (Zeiss Plan-NEOFLUOR 20x/0.5) to generate both biocytin-labeled images and DAPI-labeled images. Biocytin images were used to assess cell morphology and DAPI images were used to identify cortical layer boundaries. To further classify cell morphology, we also used a 63x lens to capture high-resolution z-stacks of biocytin filled cells, which were stitched together using ZEN software and exported as single-plane TIFF files.

### Synapse detection

Electrophysiology data was filtered through multiple quality control (QC) steps to ensure it was of good quality to detect a connection (Table S3). Additional QC criteria were applied for characterizing synapses which are discussed in the following section.

Initial data processing consisted of manual synapse detection and curve fitting (Figure 1B *Synapse Processing*) using our [Pair Analysis Tool](https://aisynphys.readthedocs.io/en/latest/interactive_tools.html#pair-analysis) (Fig S10). During recording, cells were held at −70 mV to probe excitatory connections or −55 mV to probe inhibitory connections. Within each clamp mode (voltage and current) data was binned into two ranges of recorded membrane potentials [−80, −61] (−70 mV holding potential) and [−60, −45] (−55 mV holding potential). Within each of these four groups, individual postsynaptic responses (PSC/P, Fig S10 white traces) from stimuli ≦50 Hz were aligned to the peak rate of rise of the presynaptic spike and averaged (Fig S10A blue traces). Only QC passed PSC/Ps were included in the average. Users visually identified chemical and electrical synapses from the average postsynaptic responses in each of the four quadrants and marked whether the synapse was excitatory or inhibitory. Manual connectivity calls were used to train a machine classifier^(51)^, which could reveal possible false positives or negatives that were later manually re-evaluated.

### Synapse characterization

Users manually identified the onset of detected synapses’ postsynaptic response (Fig 1B, Fig S10, yellow line). This user-defined latency was used to initialize an automated curve fit of the average response constrained to ± 100 µs of the user-defined latency (Fig S10B red and green traces). The user inspected the automated fit for a good match to the average and could refit as necessary. Once the user was satisfied with the output, the fit was manually passed (Fig S10B green) or failed (Fig S10B red). Contamination by electrical synapses or artifacts, the shape of the fit, and other factors were considered when deciding whether to pass or fail the fit. Users were also able to make notes, which were used to update and test new fitting algorithms.

Passing postsynaptic response fits were used to characterize strength, kinetics, and short-term plasticity of the synapse in a multi-stage process aimed to maximize data inclusion for each characteristic. Kinetics, rise time and decay tau, were measured as a weighted average of curve fit parameters from the two membrane potential ranges. Latency was similarly measured as a weighted average across not only membrane potentials but clamp mode as well, as we did not expect PSC/P onset to be influenced by these conditions. Latency values from the individual membrane potential/clamp mode modalities were confirmed to be within 200 µs of each other to be included in the weighted average. Thus, each synapse has a singular latency value. PSC/P amplitude was measured at a “resting state” to avoid the influence of short-term plasticity in our estimate of strength. Individual PSC/Ps recorded at the appropriate holding potential for the synapse (−55mV for inhibitory, −70mV for excitatory) and preceded by at least 8 seconds of quiescence were aligned to the presynaptic spike and averaged. This reduced average was fit with kinetic parameters initialized to those from the fit to the average of all responses. For synapse types that show short-term facilitation, the resting state amplitude is often very small and thus, we calculated a second metric to capture the near maximum strength that a synapse can produce. For this, individual PSPs were fit with parameters initialized by results of the average fit. We then calculated the 90th percentile of fit amplitudes to approximate maximum strength (this value was also used to normalize our estimate of short-term plasticity discussed below).

Short-term plasticity (STP) was measured from current-clamp PSPs in response to trains of stimuli at multiple frequencies consisting of eight pulses, followed by a variable delay, and then four more pulses (Fig 1B *Multipatch Experiment*, *Synapse Analysis*). Quantifying the magnitude of STP has taken several forms from a paired-pulse ratio (PPR, ratio of the second response to the first), to a ratio of the last pulse in a train to the first. These ratios are sensitive to noise, especially when the signal in the denominator becomes very small; thus we quantified STP using the difference between late (pulses 6-8) and initial response amplitudes, normalized by the 90th percentile response amplitude (Fig 1B), termed STP induction. STP induction was measured from the amplitude of individual PSP fits as follows (Fig 1B *Synapse Analysis*):

*(Avg(6th, 7th, 8th pulse amplitudes) - 1st pulse amplitude) / 90th percentile amplitude*

By this calculation positive values denote facilitating synapses and negative values depressing.

Recovery from STP was similarly calculated from individual PSP fits as:

*((Avg(9th-12th pulse amplitudes) - Avg(1st-4th pulse amplitudes)) / 90th percentile*

This yields positive values indicative of recovery beyond the initial state of the synapse and negative values where the synapse has not yet recovered from STP.

PSP/C variability is typically reported as the coefficient of variation (CV) of response amplitudes; however, for weak synapses the CV is dominated by noise arising from multiple factors including the release probability, quantal variance, and other biological and electrical sources^(94)^.

To access the component of the variability driven by synaptic release mechanisms, we calculated an “adjusted coefficient of variation” (aCV) that subtracts the experimental noise contribution before normalization:

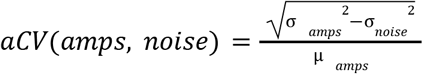

Where *μ_amps_* and *σ_amps_* are the mean and standard deviation of response amplitudes, and *σ_noise_* is the standard deviation of background noise, which is measured by performing the same amplitude measurement algorithm on regions of the recording that have no presynaptic stimulus^(61)^.

For the purposes of analyses in Figures 3 and 4, transgenic cell subclass was used to identify the pre- and postsynaptic cell type resulting in semi-layer-specificity of E-I and I-E synapses while pooling I-I synapses across all layers. Pooling was motivated by the observed homogeneity of intralaminar I-I synapses across layers (Fig S5) and allowed for more robust comparisons. Nevertheless, pooled results may contain layer-related biases due to differences in the laminar distribution of cell bodies across the inhibitory subclasses. For example, most Vip cells are located in L2/3 (Fig S5A) and thus synapses involving these cells may be biased toward upper layers^(71, 72)^.

The strength of electrical synapses (gap junctions) was quantified as a coupling coefficient and junctional conductance ^(43, 44)^. The voltage change from baseline evoked by a subthreshold long-pulse current injection (Fig 1C) was measured in both the pre- and postsynaptic cell. The coupling coefficient was measured as a least squares linear regression of the voltage change across all sweeps. For cells in which we also measured input resistance the junctional conductance was calculated as G_j_ = (1/R2) x CC/(1-CC) ^(10)^ where CC is the coupling coefficient and R2 is the input resistance of the postsynaptic cell.

### Cell characterization

#### Transgenic Expression and Cell Subclasses

Stack images of the recording site were taken in brightfield, epifluorescence (tdTomato and EGFP), and dye-filled recording pipettes. These images were filtered and overlaid on top of each other (Fig 1B *Multipatch Experiment*) to display the recording site and targeted cells. From the overlap of epifluorescence and pipette dye, we identified each cell’s transgenic expression, which was used to define its subclass. If fluorescence overlap could not be confirmed, the expression was marked as unknown.

All mouse L2/3 excitatory cells and human cells were fluorescence-negative cells and thus, morphological features were used to identify these cells as discussed below. Human excitatory cells were split into putative subclasses by layer. The inhibitory cells in human, expected to be primarily fast-spiking Pvalb cells due to their prevalence, were pooled across layers based on the consistency of inhibitory synaptic properties through the cortical depth observed in the mouse data (Fig. S6). Due to experimental constraints from tissue health and imaging challenges, sampling was primarily from the supragranular layers of cortex, and some deep subclasses were under represented. The L6 subclass was dropped from all analyses, and the L4 subclass from synapse property analyses.

#### Electrophysiology

Intrinsic characterization of individual cells was carried out similarly to that described in ^(96)^. Although we did not collect the full suite of stimuli, the long-pulse sweeps we acquired were sufficient to calculate subthreshold properties such as input resistance, sag, and rheobase; spike train properties such as f-I slope and adaptation index; and single spike properties such as upstroke-downstroke ratio, after-hyperpolarization, and width (Fig1C *Intrinsic Ephys*). For spike upstroke, downstroke, width, threshold, and ISI, ‘adaptation ratio’ features were calculated as a ratio of the spike features between the first and fifth spike. A subset of cells also had subthreshold frequency response characterized by a logarithmic chirp stimulus (sine wave with exponentially increasing frequency), for which the impedance profile was calculated and characterized by features including the peak frequency and peak ratio. Feature extraction was implemented using the IPFX python package (https://github.com/AllenInstitute/ipfx); custom code used for chirps and some high-level features will be released in a future version of IPFX.

For the human cell dataset, all electrophysiology features were aggregated and visualized using a UMAP projection ^(97)^ to gain perspective on the electrophysiological cell type (‘e-type’) distinctions present. Cells with more than 25% missing features were dropped. The remaining missing features were imputed as a distance-weighted mean of 3 nearest neighbors, and each feature was independently power transformed to a standard Gaussian. Features uninformative for known cell-type distinctions were dropped (assessed by F-score of ANOVA against layer and spininess labels), and the remaining features were visualized by UMAP projection. For the L2/3 focused analysis, the L2/3 pyramidal subclass was refined by an upper bound on input resistance of 225 MΩ, excluding L4-type cells that can overlap into L3 based on their smaller size and higher input resistance ^(24)^. These refined subclasses were visualized in the full UMAP feature space and used for the depth correlation analysis.

#### Morphology and Position

Cell morphology was qualitatively assessed from 63x maximum projection image z-stacks of biocytin filled cells and included features such as dendritic type (spiny-ness), axon origination point of inhibitory cells, and length of truncated axon (measured in pixels as a straight line from axon origination point to truncation and multiplied by image resolution to obtain distance in μm) (Fig 1C *Morphology*). Aspiny or sparsely spiny cells (inhibitory) were defined as such if their dendrites lacked or only had few protrusions. Spiny (excitatory) cells have frequent dendritic protrusions as well as an apical dendrite ^(84, 98)^. For the purposes of cell classification throughout our results, “spiny-ness” refers to this analysis. Layer 2/3 pyramidal cells in mouse were largely identified as being “spiny” as we did not have a transgenic driver for this layer. Similarly in human, all cells are identified as excitatory or inhibitory by their dendritic spiny-ness and by the presence/absence of an apical dendrite. A full list of morphological classification can be found in Table S4.

Cortical layer boundaries were determined from DAPI images, with the top of Layer 1 serving as a marker of pia and the bottom of Layer 6 as a marker of white matter. During the experiment, cell position was recorded in the fluorescent images’ reference space (Fig 1B *Multipatch Experiment*) and later coregistered with the DAPI image. Image coregistration enabled the cell soma layer to be established. Other positional metrics such as intersomatic distances (vertical and lateral), distance from pia and white matter, fractional cortical depth, and depth within the layer were also calculated from the soma position and layer boundary data using the neuron_morphology python package (https://github.com/AllenInstitute/neuron_morphology). Depth measurements were made using streamlines from the pia to WM boundaries (or the nearest layer boundary in cases where not all layers were complete in the slice). For each cell pair, the pia-WM orientation from the streamlines was averaged and used as a ‘vertical’ orientation to decompose the soma-soma separation into vertical and lateral distances.

### Connection probability estimation

We estimated the connection probability through modeling a probability distribution that depends on the experimental conditions such as the distance between the neuron pair, depth of the neurons in slices, and signal-to-noise ratio. To build a model for the connection probability, we started with a log-likelihood function of the binomial distribution, constructed its probability using multiple experimental variables, and estimated model parameters using maximum-likelihood estimation (MLE).

The log-likelihood function for the binomial distribution is defined as follows.

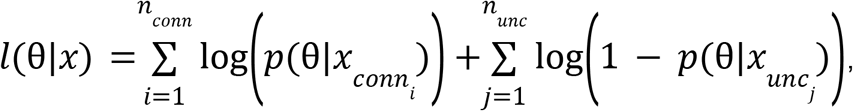

where θ is a set of model parameters, *x* is a set of experimental variables associated with each pair, *p* is the model estimate of the connection probability, *n_conn_* and *n_unc_* are the numbers of connected and unconnected pairs, *x_conn_* and *x_unc_* are subset of *x* for connected and unconnected pairs, respectively. The first sum runs over connected pairs; the second sum runs over unconnected pairs. Below, we elaborate how this probability function is constructed.

#### Gaussian model with maximum likelihood estimation

A full description of how we arrived at a Gaussian model of connection probability as a function of intersomatic distance can be found in the accompanying [notebook](https://github.com/AllenInstitute/aisynphys/blob/master/doc/connectivity_vs_distance.ipynb) and is summarized here.

Connection probability as a function of intersomatic distance of a pair was modeled as a Gaussian function centered at 0 distance:

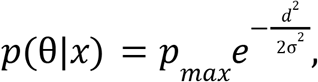

where θ = {*p_max_*, σ} and *x* = {*d*}, *p_max_* is the peak connection probability, *σ* is the distance constant of connection probability, and *d* is the lateral intersomatic distance of the pair.

The number of samples (pairs of cells) included in the model has a profound impact on our confidence in the model outputs, *p_max_* and *σ*. We used simulated distances drawn from the distribution of measured intersomatic distances and a defined *p_max_* and *σ* to assess the possible error in our fits. We found that while *p_max_* was fairly well constrained, *σ* was not, particularly for lower numbers of samples. We therefore chose to use a fixed *σ* value for groupings at the subclass level (Fig 2B), which further constrained the fit of peak connection probability. To determine the fixed *σ* value, we pooled our data from mouse into four categories based on the cell class of the presynaptic and postsynaptic cells, namely excitatory to excitatory (5,467 cell pairs), excitatory to inhibitory (2,161 pairs), inhibitory to excitatory (1,973 pairs), and inhibitory to inhibitory (6,100 pairs). This allowed us to have several thousand samples in each group to obtain a better estimate of *σ* (Fig 2A). This showed a trend for shorter *σ* values for within class connections and longer *σ* values for across class. In order to more fully determine if the *σ* of these four groups could have been drawn from the same distribution we simulated 10,000 experiments with a true *σ* that varied between those measured from experimental data. We then did a pair-wise comparison of the *σ* ratio for each unique comparison among the four groups and calculated the percentile of this distribution where the measured ratio fell. When correcting for multiple comparisons we found that while the I→I and I→E connectivity profiles likely have *σ* that are not drawn from the same distribution, we could not rule that out for the other comparisons. This analysis was also conducted for a comparison of chemical versus electrical connections among inhibitory cells, with 1,000 simulated trials. From this analysis we chose to fix *σ* for individual matrix elements of chemical synapses (Fig 2B) to 95 μm for within class and 125 μm for across class, and to fix *σ* for electrical synapses (Fig S4A) to 77 μm. A similar procedure was conducted for human data, resulting in a fixed *σ* of 140 μm (Fig 5A).

#### A unified model of connection probability adjustment

We extended the analysis of the connection probability as a function of intersomatic distance to include the effect of tissue slicing and false negative detection of connections due to signal-to-noise ratio. We created a unified model that applies these adjustments to the connection probability, as a function of the pair distance (discussed above), the presynaptic axon length, the depth of the cells, and the detection power. Using the same log-likelihood function described above, we extended the probability to the following.

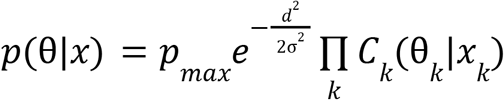

where θ = {*p_max_*, σ, ρ, µ*_ax_*, σ*_depth_*, µ*_depth_*, σ*_det_*}, *x* = {*d*, *l_ax_*, *z_depth_*, *p_det_*}, and product over k runs over the following three corrections factors. The correction functions *C_k_*, their parameters θ*_k_*, and their variables *x_k_* are described below.

The model for the presynaptic axon length is a binary step function with a threshold at 200 µm as axons were not measured past this point (Table S4). Also, because the number of neurons with axons measured less than 200 µm were few, we did not have sufficient data to determine the function shape below this threshold. Therefore, we used a single adjustment ratio for the correction.

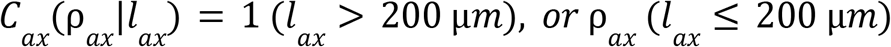

The model for the average depth of the pair of neurons is as follows.

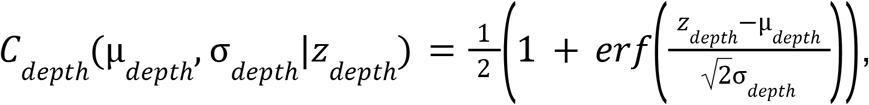

where *erf* is an error function:

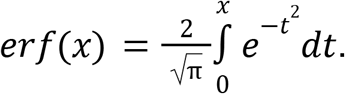

The detection power (that is proportional to signal-to-noise ratio) is defined as 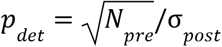, where *N_pre_* is the number of presynaptic test spikes and σ*_post_* is the RMS noise-level of the postsynaptic neuron. The model for the detection power *p_det_* is also an error function, but in a log space, because the synaptic weight distribution is expected to be a log-normal distribution.

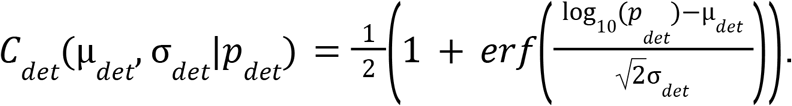

We did not see a saturation of the connection probability when the presynaptic cell was inhibitory (Fig S1D), suggesting that there are potentially a large number of undetected synapses. However, we did not want to overestimate the connection probability in the range of detection power where we did not have sufficient data. Therefore, we applied the following constraint to the fit.

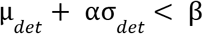

Where α ∼ 0. 6745, specifying the quartile of the integrand Gaussian of the error function, and β ∼ 4. 6613, specifying the quartile of the detection power distribution in our data. This constraint ensures saturation in the high detection power region where data are scarce.

The models and the parameters for these three corrections are determined individually for each variable (Fig S1B-D) and incorporated into the likelihood function used for the distance adjustment (Fig S1A). When we estimated final *p_max_*, we performed a single-parameter MLE, fixing all the other model parameters to pre-determined values. Four models were used for mouse connectivity reflecting the varying effects that each variable has on excitatory versus inhibitory and within versus across class. Matrix elements at the subclass level (Fig 2B, S2A) were determined to be one of E→E, E→I, I→E, or I→I and the appropriate model was applied to adjust *p_max_* (Fig S1F).

In the case of human synapses, the data was insufficient to fully constrain the complete connection probability adjustment model, and we applied an adjustment for lateral intersomatic distance only. However, calculations suggested that the remaining adjustments are likely much smaller than in the mouse dataset (<50%) due to higher detection power arising from lower noise and more recorded spikes.

#### Estimating connection probability and confidence intervals

The goal of optimizing the Gaussian MLE was to compare connection probabilities across different cell groups that may have been sampled at different intersomatic distances and have variable presynaptic axon lengths and detection power. Thus, in addition to fitting *p_max_* we wanted to calculate a confidence interval of connection probability. Our initial approach was to analyze hundreds of resampled iterations of data from each matrix element. However, this method is computationally expensive and starts to break down when connection probability or the number of samples is very low. We determined the confidence intervals based on the log-likelihood function ^(99)^, assuming our log-likelihood function is asymptotically proportional to the χ^2^ -distribution (− 2*l*(*p_max_*|*x*) ∼ χ^2^ (*p_max_*); Wilks’ theorem^(100)^). Namely, we estimated upper and lower bounds of the CI as *p*_*max,CI*_ such that

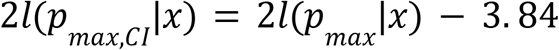

Where 3.84 is 95-percentile of the χ^2^-distribution with one degree of freedom. The computation of the CIs are done by MINOS algorithm in iminuit package^(101)^. The estimated confidence intervals were used to shade connection probability heatmaps in Figure 2B, 5A, and Figure S2A.

The parameter optimization can result in values for *p_max_* (or CI bounds) greater than 1 in cases where the data is not well fit by the fixed-width Gaussian, typically because a class of connections has either low sampling or a true underlying connection probability function with a distinct shape or size. In the resulting figures (Fig 2B, 5A, S2A), *p_max_* values and CI upper bounds were clipped at 1, reflecting the fact that such data should be better fit by a flat-topped curve that approaches 1 at its maximum (Fig. S2B). The unclipped values are available in our data and code release for applications like modeling, where estimating the true connection probability vs. distance function may be more important than interpretation of *p_max_* as a probability.

### Modeling synapse behavior

#### Stochastic quantal release model

We developed a model of stochastic vesicle release and synaptic dynamics that expands upon standard models ^(102)^ and is similar to some recent models ^(80, 81, 103)^. The model was designed to meet several criteria. First, it should give estimates of synaptic quantal parameters (number of release sites, probability of release, and quantal size) and dynamic parameters (facilitation, docking, etc.). The quantal parameters alone allow the model to predict the overall distribution of response amplitudes (Fig S11A). By allowing quantal parameters to change in response to the history of activity, the model is able to account for changes in the response distribution (Fig S11B,C). Next, the model should operate on individual spike times and response amplitudes rather than requiring structured or repeated stimuli. This ensures that the model can access the complete distribution of response amplitudes and any correlations between adjacent responses. Additionally, operating on individual events rather than averages ensures that we use all available data for each synapse, regardless of the applied stimuli. Finally, the model should fail gracefully in cases with low signal-to-noise ratio by indicating low confidence over its parameters rather than returning unreliable values.

We begin with a standard quantal model with three parameters: *N*, the number of release sites; *q*, the amplitude of the postsynaptic response to a single vesicle, and *P_r_*, the probability that each release site will release a vesicle in response to one action potential. We make simplifying assumptions that all release sites in a connection share the same values of *P_r_* and *q*, and that the response to multiple vesicles released simultaneously is simply the linear sum of individual responses. This component of the model simply predicts that the number of vesicles released per spike is defined by a binomial distribution with parameters *N* and *P_r_*, and that the distribution of response amplitudes is the same with an additional scaling factor *q*.

The measured response amplitudes in our dataset only occasionally show binomial characteristics; however in most cases, background recording noise and quantal variability obscure the underlying discrete distribution. We model these sources of variability as independent Gaussian distributions for measurement noise and quantal variance, with parameters σ_m_ and σ_q_, respectively. These combine with the binomial distribution to make a weighted Gaussian mixture model that, on its own, does a decent job of approximating the overall distributions of event amplitudes. The model response amplitude probability distribution *P(X; θ)* is calculated as:

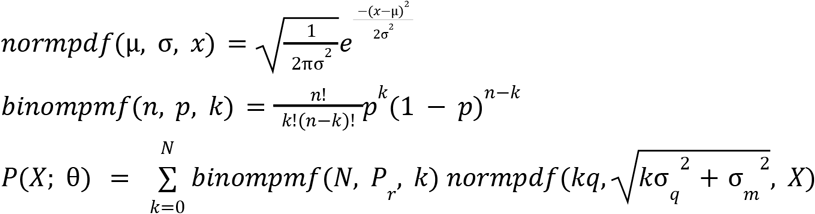

#### Short term plasticity

Synapses undergo short term plasticity, which we model as changes in the expected distribution of response amplitudes over time ^(102)^. Each incoming action potential causes an instantaneous modification to the quantal parameters *N* and *P_r_*, which then recover back to their initial values by exponential decay until the time of the next action potential. Although we only seek a phenomenological description of the synapse, this description is designed to mimic three major classes of synaptic dynamics: vesicle depletion, facilitation, and calcium channel inactivation. Vesicle depletion is implemented by using the response amplitudes and *q* to estimate the most likely number of released vesicles following each spike, which is then subtracted from the releasable vesicle pool. Recovery from vesicle depletion is modeled as an exponential with time constant *τ_r_*. Facilitation and calcium channel inactivation increase or decrease the release probability, respectively, for every incoming spike, and decay back to the initial release probability with time constants *τ_f_* and *τ_i_*. In addition to the decay time constants, these mechanisms introduce an extra two parameters: the amount of facilitation *a_f_* and inactivation *a_i_* per spike.The complete algorithm then looks like:

1. Initialize state variables:

a. N_j_ = N_r_
b. *depression = 0*
c. *facilitation = 0*
2. For each spike *j* at time *t_j_*:

a. Let *dt = t_j_ - t_j-1_*
b. Recover state variables:

i. N_j_ += (N_r_ - N_j_) * (1 - e^−dt/τD^)
ii. *depression *= e^−dt/τD^*
iii. *facilitation *= e^−dt/τF^*
c. Let *P_j_ = (1 - depression) * (P_r_ + (1 - P_r_) * facilitation)*
d. Define distribution parameters *θ = {N_j_, P_j_, q, σ_q_, σ_m_}*
e. Estimate likelihood of measured response amplitude *P(Amp_j_ | θ)* or generate a random sample drawn from *P(θ)*
f. Apply post-spike modifications to state variables:

i. *vesicle_pool -= Amp_j_ / q*
ii. *depression += a_D_ * (1 - depression)*
iii. *facilitation += a_F_ * (1 - facilitation)*

Occasionally a presynaptic stimulus is not followed by a detectable spike, which could be caused by a genuine spike failure or simply by a failure of the spike detection algorithm. In these cases, we assume that a spike did occur for the purpose of updating the model state variables, but we incur a short timeout during which we stop accumulating evidence toward the overall model likelihood.

For any combination of parameters, we estimate a *likelihood* that a recorded set of amplitudes *A_1..m_* could be generated by the model by the mean log probability density for each response:

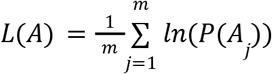

If most response amplitudes fall within the modeled regions of high probability density, then the total likelihood for that parameter set will be high. Low-likelihood models result from either bad parameters (where response amplitudes fall in low density regions of the model probability distribution) or from insufficiently selective parameters (where the probability distribution is spread out over too much area).

#### Parameter search and optimization

With a measure of model likelihood, we can now attempt to find a set of parameters that maximize this value, yielding a model that best explains the data. Most prior methods use a metric similar to the likelihood defined above along with an optimization method to efficiently find a single point in the parameter space that is most consistent with the recorded data. However, minimization is notoriously difficult in this domain because the model parameters are underconstrained--there exist many solutions that adequately explain the recorded data, and thus large differences in the optimal parameters may simply result from noise or experimental artifacts, rather than physiological differences between synapses ^(103, 104)^. To avoid this outcome, we measure the model performance at every point in a large parameter space, thereby identifying the region of the parameter space consistent with the responses recorded from each synapse (Figure S11D). This is similar to some recent methods ^(80, 81)^, but differs in that we have implemented a simpler (and thus less computationally expensive) model in order to test many more parameter combinations uniformly.

All combinations of the parameters in a 7-dimensional space were tested, for a total of 6.2 M model tests per synapse. At every point in this parameter space, the quantal amplitude *q* was estimated and optimized using a minimization algorithm (scikit.optimze). Although this minimization strategy fails to find reliable optima when operating over several parameters, we found it to be reliable in the context of this simpler single-variable optimization. The resulting records for each synapse thus include measured model likelihood as well as the optimal value of *q* at each point in the parameter space. This yields an 8 dimensional image that serves as a “fingerprint” describing the unique dynamics of each synapse. The size of the parameter space and the number of synapses in our dataset together make this a computationally expensive operation. To reduce this cost, we optimized the core routines used in the model using the python packages numpy and numba.

### Synapse typing

To visualize the relationship between cell subclass and synapse properties, we used the UMAP dimensionality reduction method ^(97)^ to organize our synapses into a 2 dimensional space for visualization. For each synapse, the vesicle release model described above was run on a large parameter space, yielding 6.2 million “features” that collectively describe the regions of parameter space that are (and the regions that are not) compatible with the recorded data. We then used sparse PCA to reduce this down to 50 features. This set of features provides a fingerprint of any information available to the release model, including synaptic strength, stochasticity, and short-term plasticity. Finally, all features were normalized (scikit-learn), then passed to UMAP for the final dimensionality reduction. The resulting 2D space could then be visualized alongside other synaptic and cellular features to investigate the structure inherent in the data. To verify this structure is not an artifact of the reduction to two dimensions, we repeated the analysis for three dimensions but found a similar 2D structure flattened in 3D space.

---

**Figure Supplement 1.**
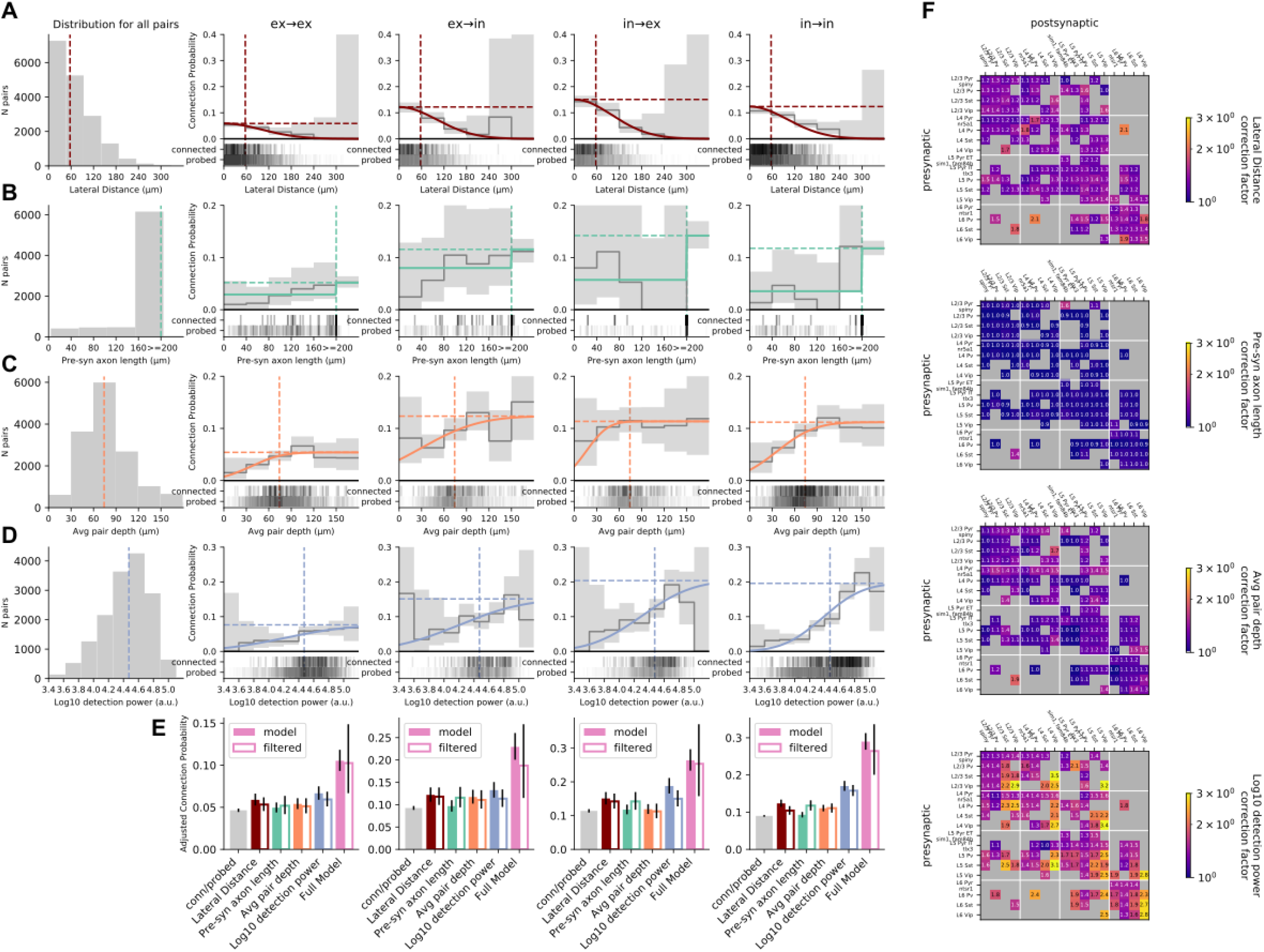
Connectivity Adjustments. **A**. From left: Distribution of lateral intersomatic distance (vertical dotted line denotes median throughout row). Connection probability as a function of intersomatic distance for E→E, E→I, I→E, and I→I pairs and 95% confidence interval (grey line/shading) with thresholded fit in the colored line. Fit for the relationship between connection probability and intersomatic distance was a Gaussian. Horizontal dotted line denotes fit *p_max_*. Raster below plot shows intersomatic distance of pairs that were probed for connectivity along with pairs that were connected. **B**. Same plots as A for presynaptic axon length measured from biocytin fills. If the axon was measured to at least 200 μm the axon was not measured further except in rare occasions. Fit for the relationship between connection probability and presynaptic axon length was a step function at 200 μm. **C**. Same plots as A for the average depth of the cell pair from the slice surface. In this case the relationship between connection probability and average pair depth is fit with an error function. **D.** Same plots as A for detection power. Detection power combines the signal to noise ratio of the postsynaptic cell with the number of spikes elicited by the presynaptic cell to probe the connection (see Methods). Detection power as a function of connection probability was also fit with an error function. **E**. Comparison of model fit *p_max_* (solid bar) to *p_max_* of data filtered above the median (open bar; vertical dashed lines in A-D; in the case of intersomatic distance inclusion was for distances shorter than the median) for each feature in A-D compared to raw connection probability (connected / probed). This highlights the overall effect that each feature has on peak connection probability. The pink bars show *p_max_* for the full model (see Methods). Error bars denote 95% confidence interval. **F**. Adjustment factor of each feature applied to each element in the matrix in Fig 2B.

**Figure Supplement 2.**
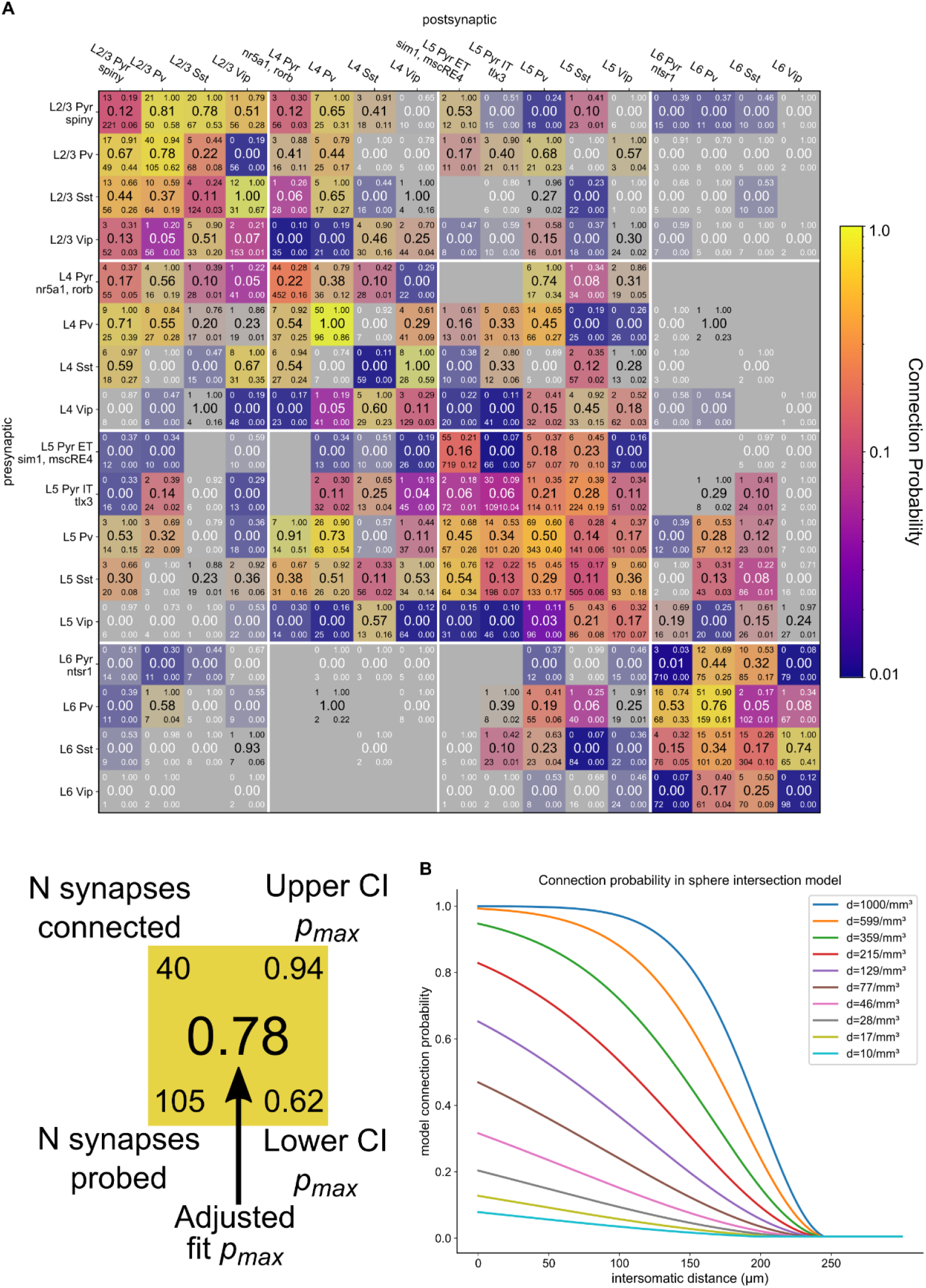
Connectivity Matrix. **A.** The connectivity matrix in Fig 2B with additional details highlighted in the expanded element below the matrix. The center number is the fully adjusted (see Figure S1) *p_max_*. The upper and lower numbers on the left of each element are the number of connections found (upper) and number of connections probed (lower). The numbers on the right of each element are the upper and lower 95% confidence interval, respectively. **B**. Probability of connection versus intersomatic distance in a simplified model that assumes a constant density of synapses inside the volume intersection of two spheres. Changing the density of synapses can result in profiles that look qualitatively like an exponential decay, a Gaussian, or a sigmoid.

**Figure Supplement 3.**
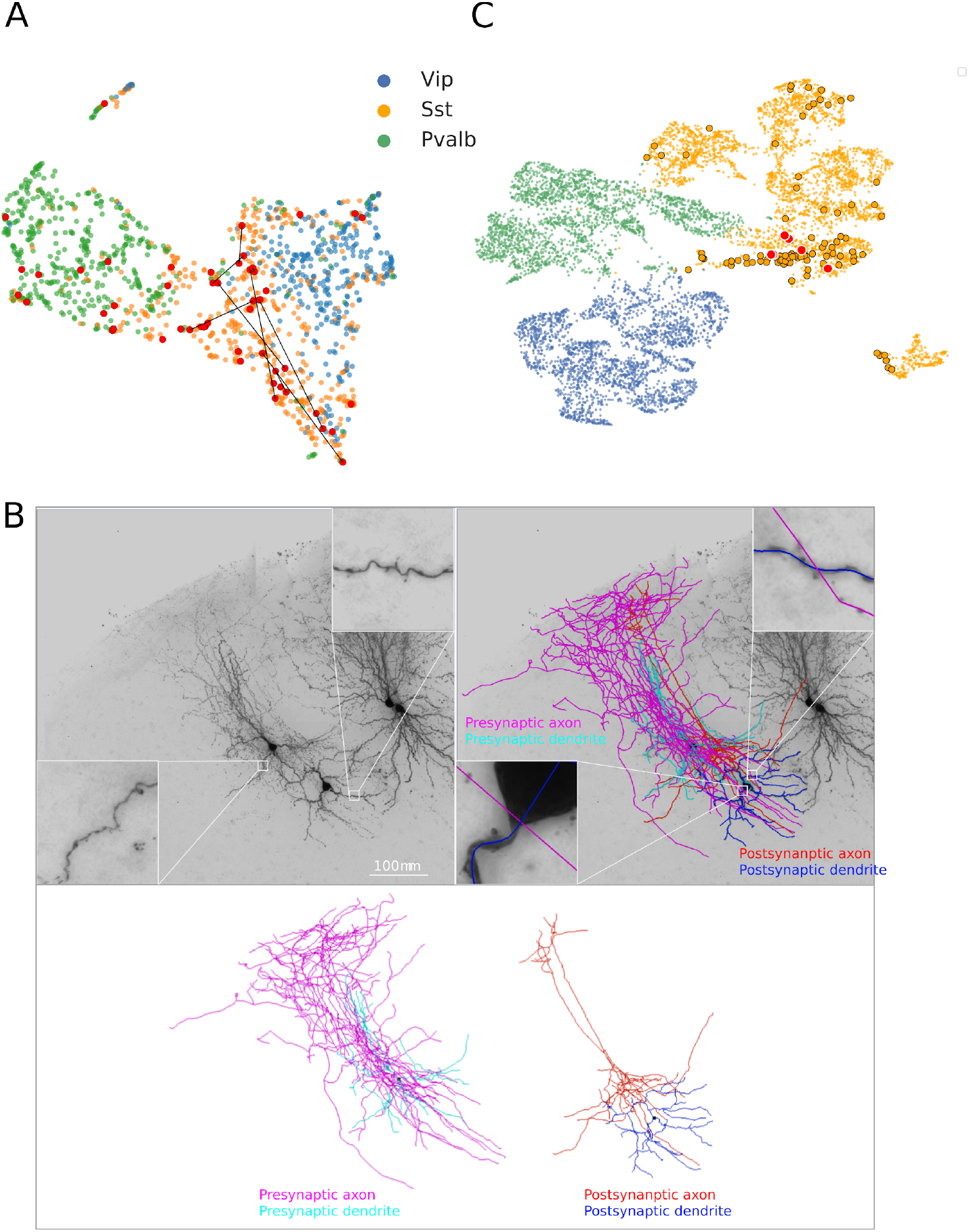
Intrinsic uMap and morphologies of Sst recurrent connections. **A**.UMAP projection of electrophysiology feature space of all mouse inhibitory interneurons. Sst-cre/flp to Sst-cre/flp chemical synaptic connections (colored lines) are overlaid. Umap is color coded by the transgenic cell class. 9 out of 16 Sst to Sst connections had pre- and post-synaptic neurons that mapped with other Sst neurons and apart from Pvalb-cre/flp neurons. Only Sst to Sst connections that fall within the Sst island are indicated with a line. Sst-cre/flp cells that are part of a connection are indicated in red. **B**. Biocytin image of connected Sst-cre to Sst-cre connected cells with Sst-like morphologies (left). Insets show sparsely spiny dendrites for pre and post synaptic cell. Biocytin image with overlaid reconstructions (right). Insets show sites of close apposition between axon and dendrite of connected neurons. Reconstructions shown in C, show independently (bottom). **C**. UMAP projection of IVSCC patchseq, FACS, and mSeq feature space of mouse inhibitory interneurons. Large dots in the Sst space highlight those from mSeq with cells from Sst→Sst connections further highlighted in red.

**Figure Supplement 4.**
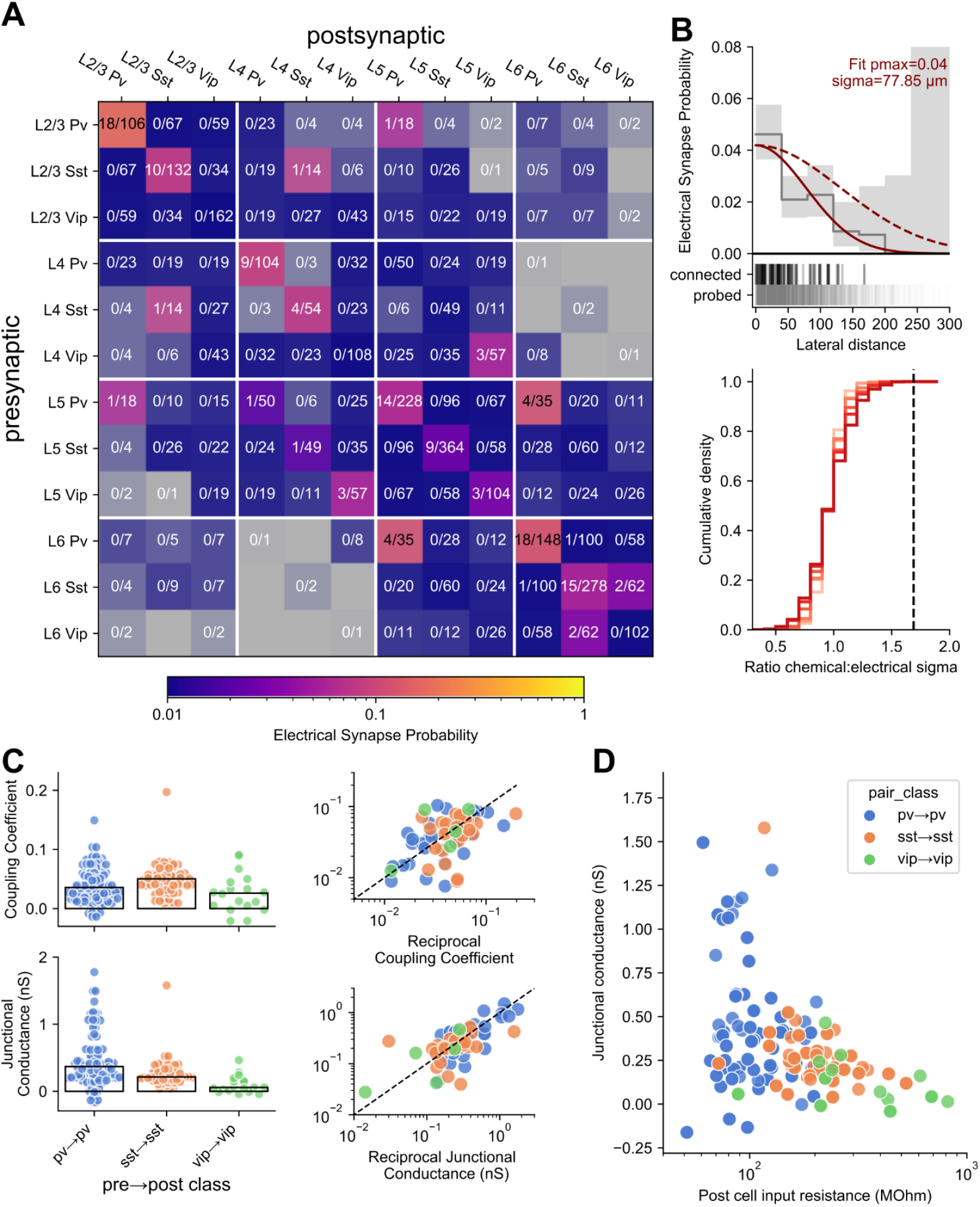
Electrical Synapses. **A.** Electrical connectivity matrix among inhibitory cells in each layer from mouse. **B**. (upper) Electrical connection probability as a function of lateral intersomatic distance with 95% confidence interval (grey line/shading) fit with a Gaussian (solid red line). Dotted red line shows Guassian fit of chemical I→I connections (normalized to electrical *p_max_*) for reference and to highlight shorter *σ* of electrical connections. (lower) Cumulative histogram of *σ* ratio comparing chemical and electrical connections. 1000 experiments were simulated in which the true *σ* for electrical and chemical connections was set to six evenly distributed values between 65 and 140 μm (light to dark red). We then measured the ratio of the Gaussian fit *σ* between chemical and electrical connections from those 1000 experiments which are plotted here as a cumulative histogram (a value of 1 indicates that the Gaussian profile of electrical and chemical connections has the same *σ*). The dotted vertical line denotes the measured *σ* ratio between chemical and electrical synapses and sits beyond the 99th percentile for all simulations. **C**. Coupling coefficient (top) and junctional conductance (bottom) of recurrent I→I electrical connections. Left plots show a scatter where each dot is the value for a single unidirectional electrical connection and bars denote the median. Right plots show coupling coefficient and junctional conductance of each electrical connection vs it’s reciprocal connection (dotted line is unity line). **D**. Junctional conductance as a function of input resistance of the postsynaptic cell.

**Figure Supplement 5.**
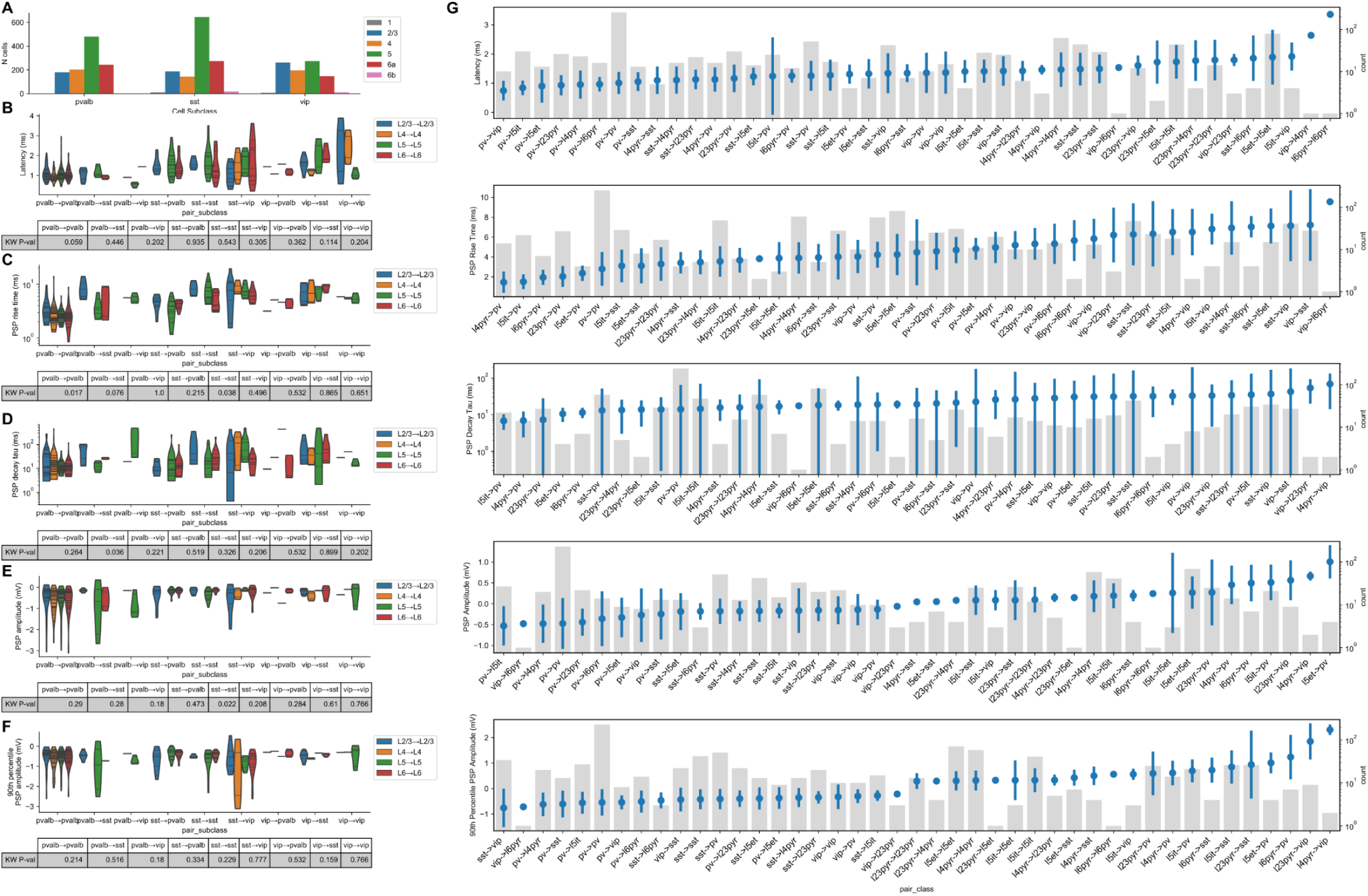
Strength and Kinetics. **A**. The distribution of inhibitory cells according to layer. **B**. Latency of inhibitory → inhibitory connections for all of the combinations among Pvalb, Sst, and Vip. For each connection element they are grouped by layer to estimate the variance of I → I latency across layer. The table below shows the p-value from a Kruskal-Wallis (KW) test which suggests that within layer I → I latency does not vary across layers. **C-F**. The same as B for PSP rise time (**C**), PSP decay tau (**D**), PSP resting state amplitude (**E**), and PSP 90th percentile amplitude (**F**). **G**. Average strength or kinetic measurement of each element in the 8 x 8 matrix from Figure 3 with standard deviation sorted from lowest to highest (blue dots, left axis) and number of pairs within each element shown in the grey bars (right axis).

**Figure Supplement 6.**
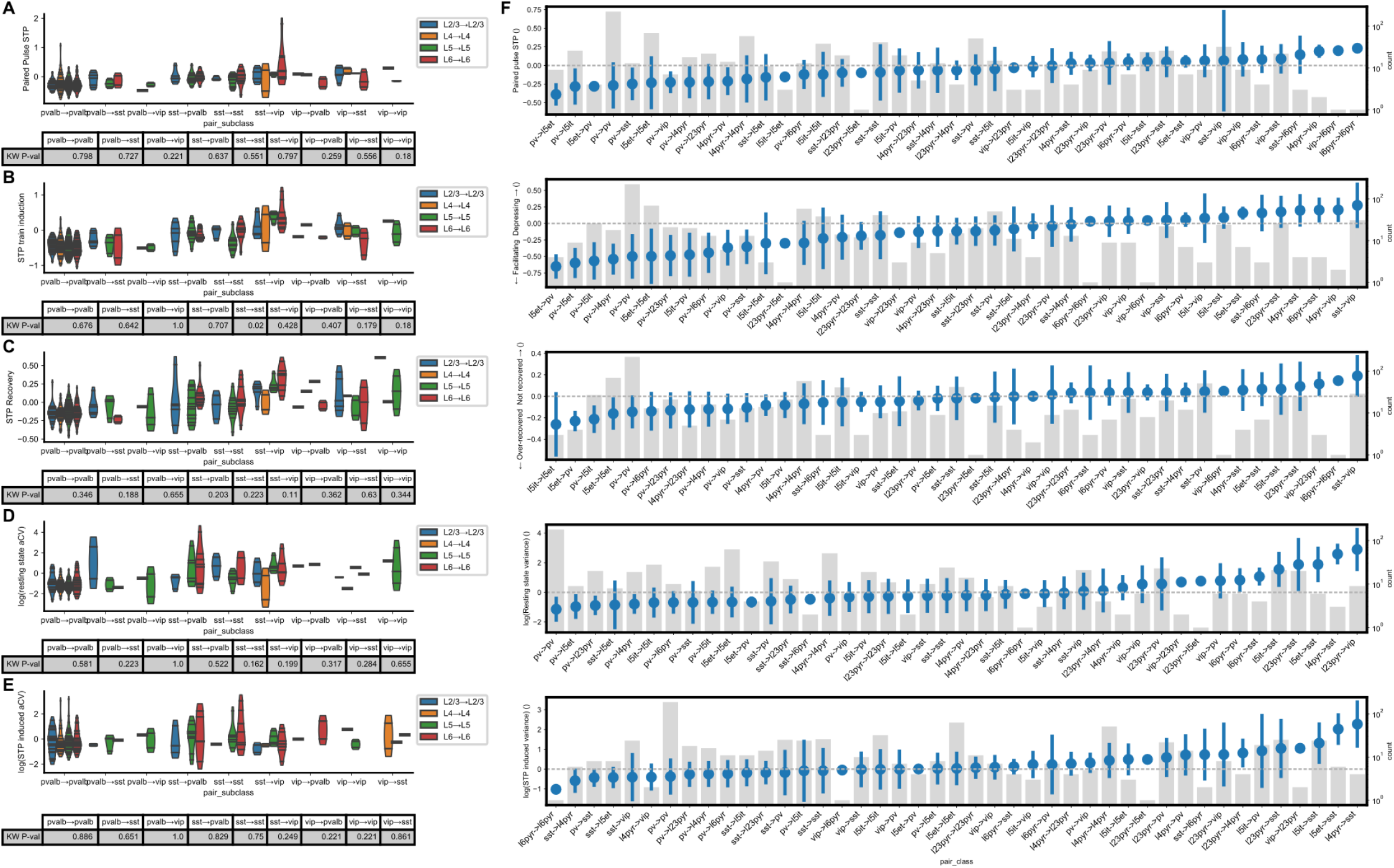
Dynamics. **A**. Paired pulse ratio (stp_initial_50hz) of inhibitory → inhibitory connections for all of the combinations among Pvalb, Sst, and Vip. For each connection element they are grouped by layer to estimate the variance of I → I PPR across layer. The table below shows the p-value from a Kruskal-Wallis (KW) test which suggests that within layer I → I PPR does not vary across layers. **B-E**. The same as A for STP induction (**B**), STP recovery (**C**), resting state variability (**D**), and STP induced variability (**E**). **F**. Average dynamics measurement of each element in the 8 x 8 matrix from Figure 4 with standard deviation sorted from lowest to highest (blue dots, left axis) and number of pairs within each element shown in the grey bars (right axis)

**Figure Supplement 7.**
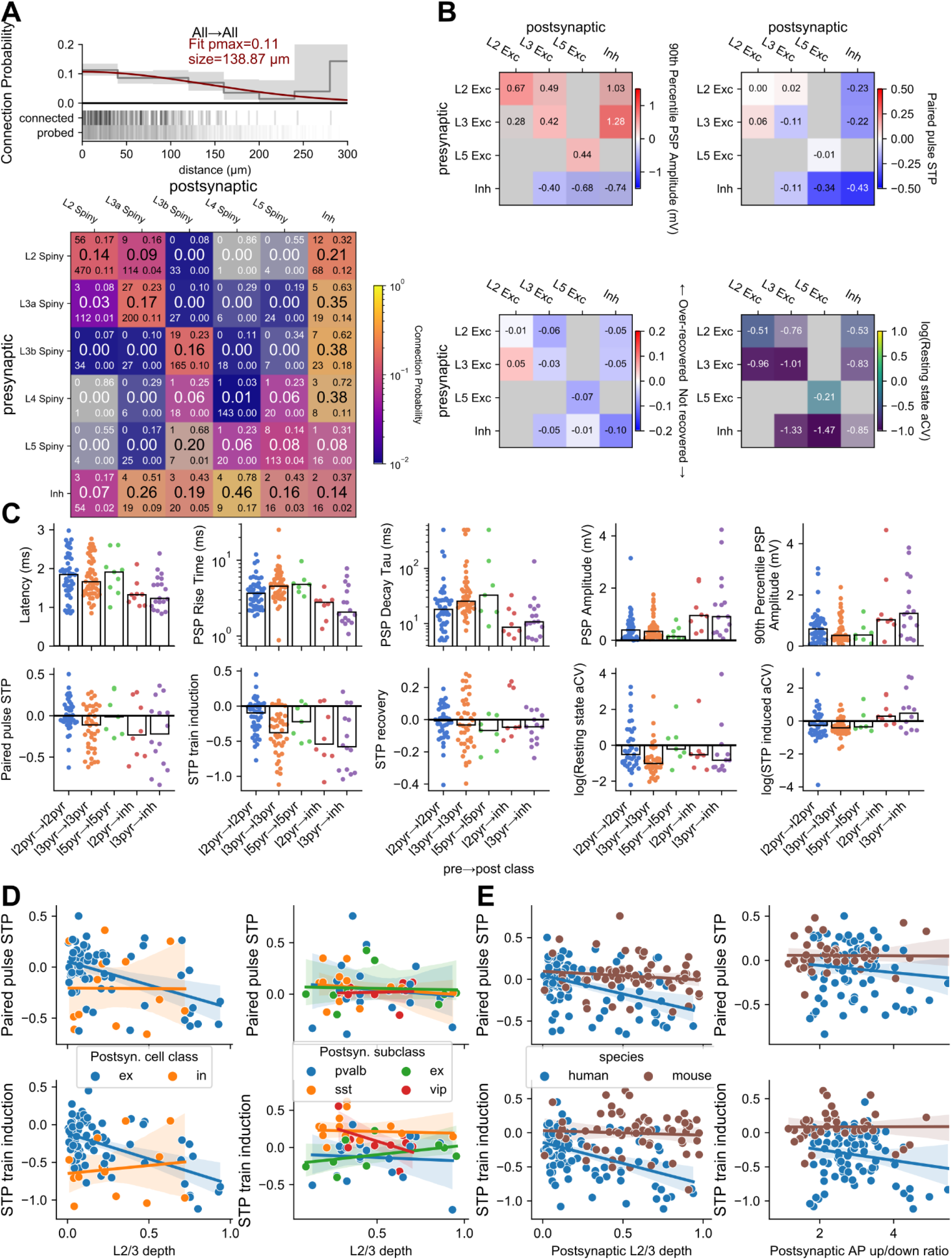
Human Data: **A.** Gaussian fit of connection probability for all human synapses. Connection probability as a function of lateral intersomatic distance was fit with a Gaussian (red line). Output parameters pmax and size describe the max connection probability and sigma of the Gaussian. Grey line and area are 40um binned average connection probability and 95% confidence interval. Raster below shows distance distribution of connections probed (bottom) and found (top). Bottom panel shows connectivity matrix from Fig 5A with additional details quantified, as in Fig S2A (number tested/probed, left; lower/upper 95% CI bounds, right). **B.** Additional STP and variability metrics of human synapse. Matrices are organized by layer for excitatory cells, with inhibitory cells grouped across layer. Each element is colorized by the grand average (text in each element) according to the colormap with the saturation scaled to the standard error. Two or more pairs were required to fill in an element. **C.** Summary plots for a range of synapse strength, timing, STP, and variability measurements. Each dot corresponds to the average response from one synapse. Responses are shown for a subset of matrix elements with sufficient sampling. **D-E.** Additional correlates of STP variability in synapses from L2/3 pyramidal cells. D shows dependence on postsynaptic cell class/subclass in human (left) and mouse (right). E shows decreased correlation with depth (left) and AP upstroke/downstroke ratio (right) when indexed to the postsynaptic cell.

**Figure Supplement 8.**
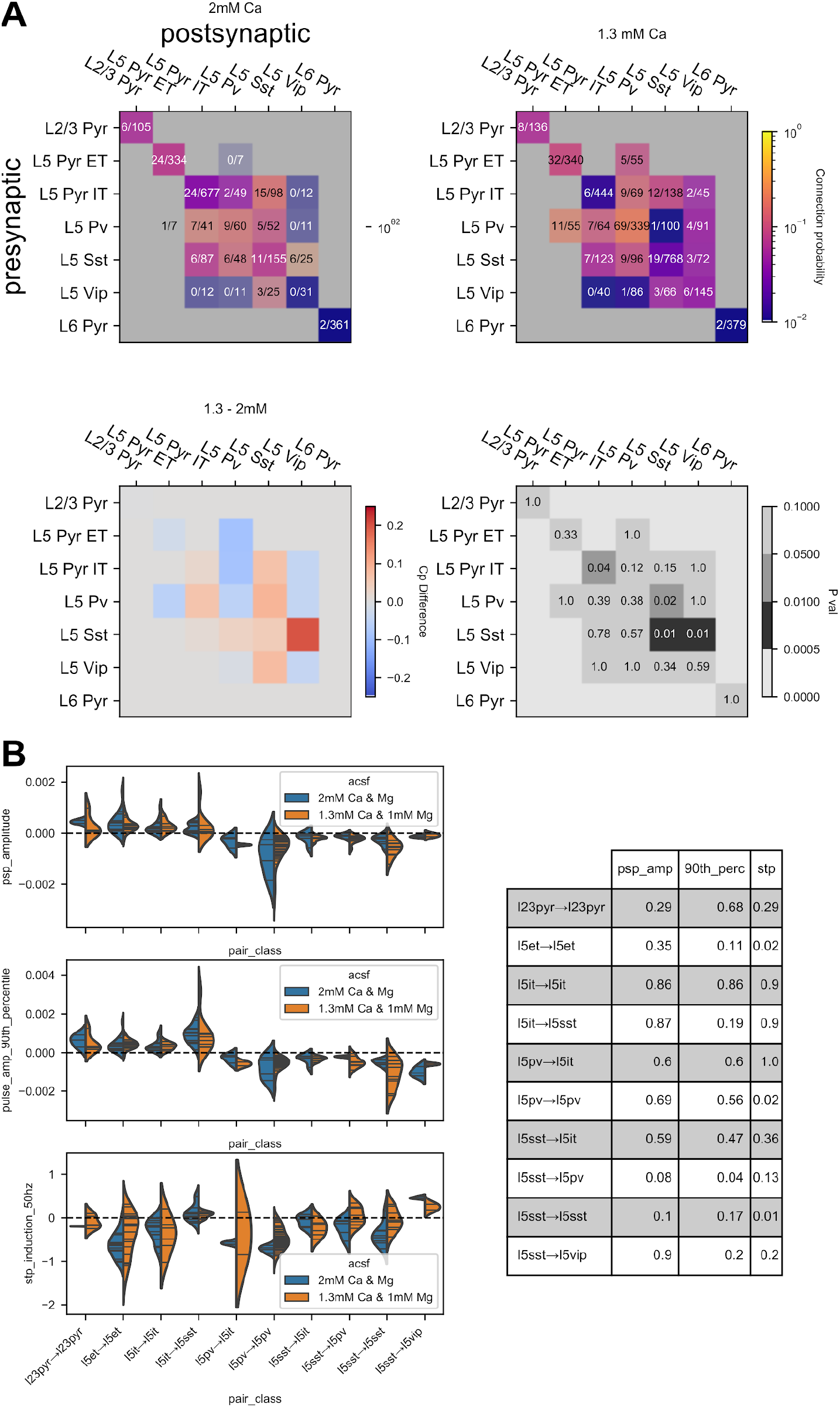
External Calcium Concentration. **A.** Connectivity matrices of elements in mouse which were probed with both 2 mM external calcium and 1.3 mM external calcium (top row). The bottom left matrix is a difference of the 2 mM matrix from the 1.3 mM matrix. Red elements are those that showed higher connectivity in 1.3 mM calcium and blue elements those that showed higher connectivity in 2 mM calcium. The bottom right matrix shows uncorrected p-values from a Fisher-exact test for each element. **B.** Violin plot of measured PSP amplitude (resting state and 90th percentile) and induced STP or pairs for each element in 2 mM external calcium (blue) and 1.3 mM external calcium (orange). Measurements from individual pairs denoted by black lines within violin. Pairs/measurements are from different experiments, Kolmogorov-Smirnov p-values are shown in the table to the left for each element.

**Figure Supplement 9.**
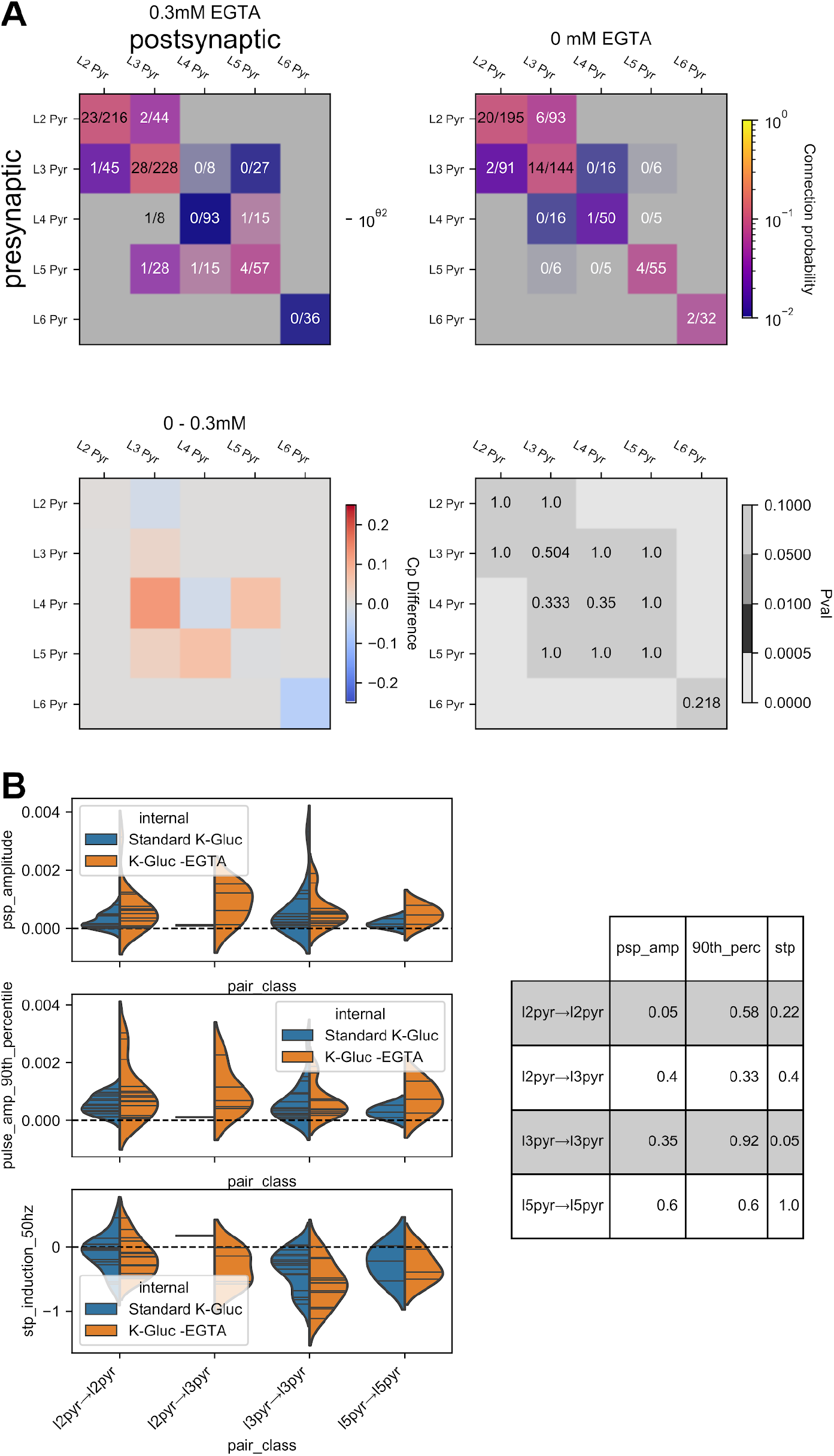
Internal EGTA Concentration. **A.** Connectivity matrices of elements in human which were probed with both 0.3 mM internal EGTA and 0 mM internal EGTA (top row). The bottom left matrix is a difference of the 0.3 mM matrix from the 0 mM matrix. Red elements are those that showed higher connectivity in 0 mM calcium and blue elements those that showed higher connectivity in 0.3 mM calcium. The bottom right matrix shows uncorrected p-values from a Fisher-exact test for each element. **B.** Violin plot of measured PSP amplitude (resting state and 90th percentile) and induced STP or pairs for each element in 0.3 mM internal EGTA (blue) and 0 mM internal EGTA (orange). Measurements from individual pairs denoted by black lines within violin. Pairs/measurements are from different experiments, Kolmogorov-Smirnov p-values are shown in the table to the left for each element.

**Figure Supplement 10.**
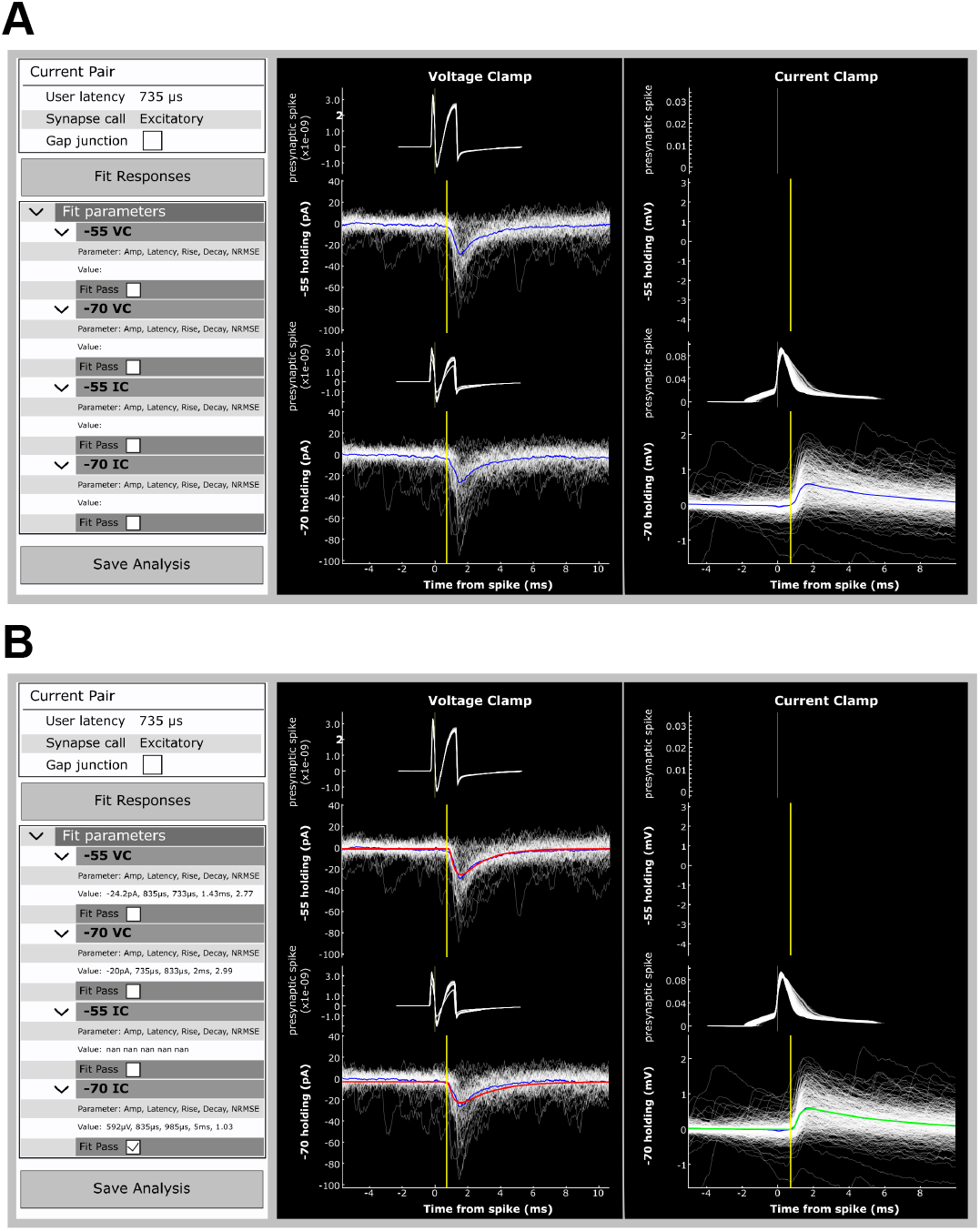
Pair Analysis Tool. **A.** Pair analysis tool used to analyze synapses. Postsynaptic responses and presynaptic spikes (white traces) from each pair were divided by clamp mode (voltage clamp on the left, current clamp on the right) and then again by baseline potential (depolarized potentials on the top, hyperpolarized potentials on the bottom). Responses from each quadrant were averaged (blue trace). If a synapse was identified the user would select the type (excitatory or inhibitory) from the menu on the left and move the yellow line in any quadrant to the onset of the response (all lines are linked). The user would then select “Fit Response”. **B.** The fitting algorithm produces a fit of the PSC/P and plots in red (QC fail, NRMSE too high) or green (QC pass). Fit parameters for each quadrant are printed in the left menu. Users could shift the yellow line and refit to obtain a better fit. When the user was satisfied with the fit result or could not obtain a passing fit the analysis was saved.

**Figure Supplement 11:**
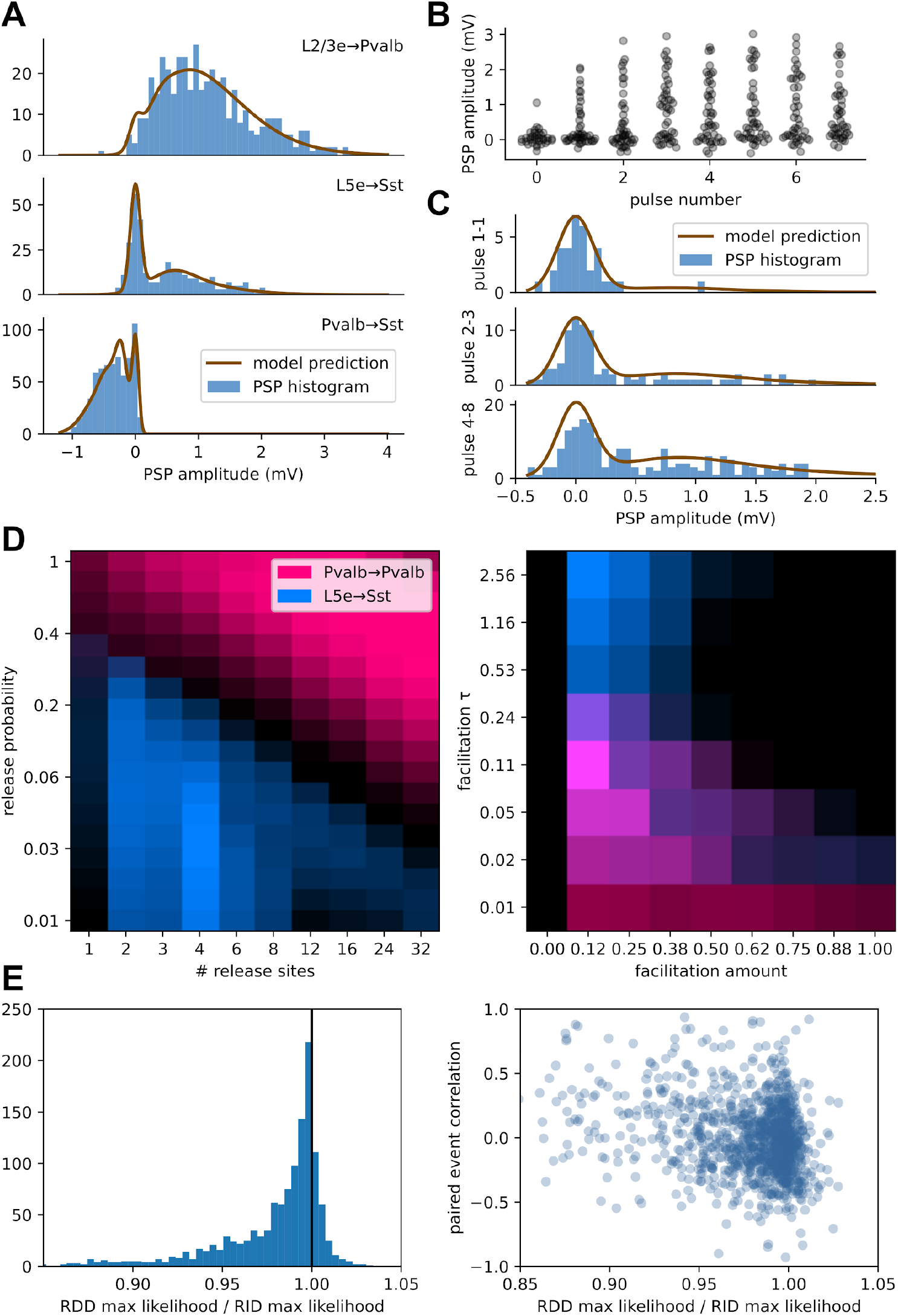
A model of quantal release and short term plasticity. **A.** PSP amplitude histograms for three synapses with the model’s average predicted amplitude distribution overlaid. **B.** PSP amplitudes facilitating and reduction of synaptic failures across a 50 Hz spike train for the example L5e→Sst synapse in (A). **C.** Histograms showing PSP amplitudes for the same synapse with model distribution predictions overlaid, showing history-dependent adjustment in model state. **D.** Estimates of model likelihood (bright colors are higher likelihood) across a range of parameters for two synapses. Each image is a maximum projection across all other axes in the model parameter space. **E.** Model results support release-independent depression mechanisms. Left: histogram of the ratios between release-dependent max likelihood and release-independent max likelihood, showing an overall preference for release-independent model parameters. Right: comparison of the same RDD/RID ratio to paired event correlations, with little overall effect. Release-dependent depression should result in negative paired event correlations.

**Figure Supplement 12:**
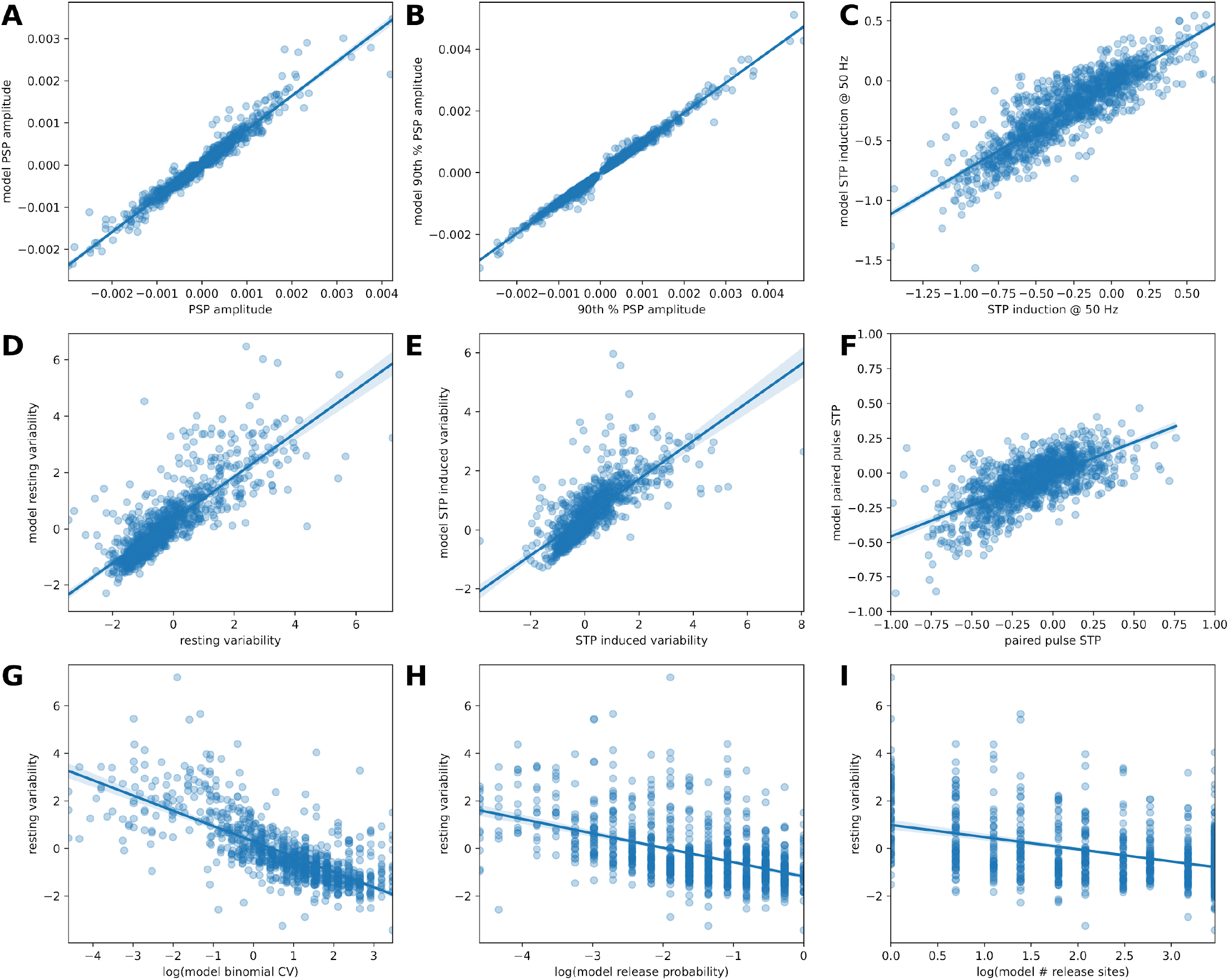
Comparison between model behavior and measured synapse features. For each synapse, the maximum likelihood model parameters were used to simulate experimental data. Measurements were then performed on the simulated response amplitudes and compared to identical measurements from the recorded data. **A-B.** Measures of synaptic strength correlate strongly with model results. **C-F.** Two measures of STP and variability. **G.** Resting state variability correlates with the binomial CV derived from model parameters. **H-I.** Release probability is more strongly correlated with variability than number of release sites.

**Table Supplement 1.**
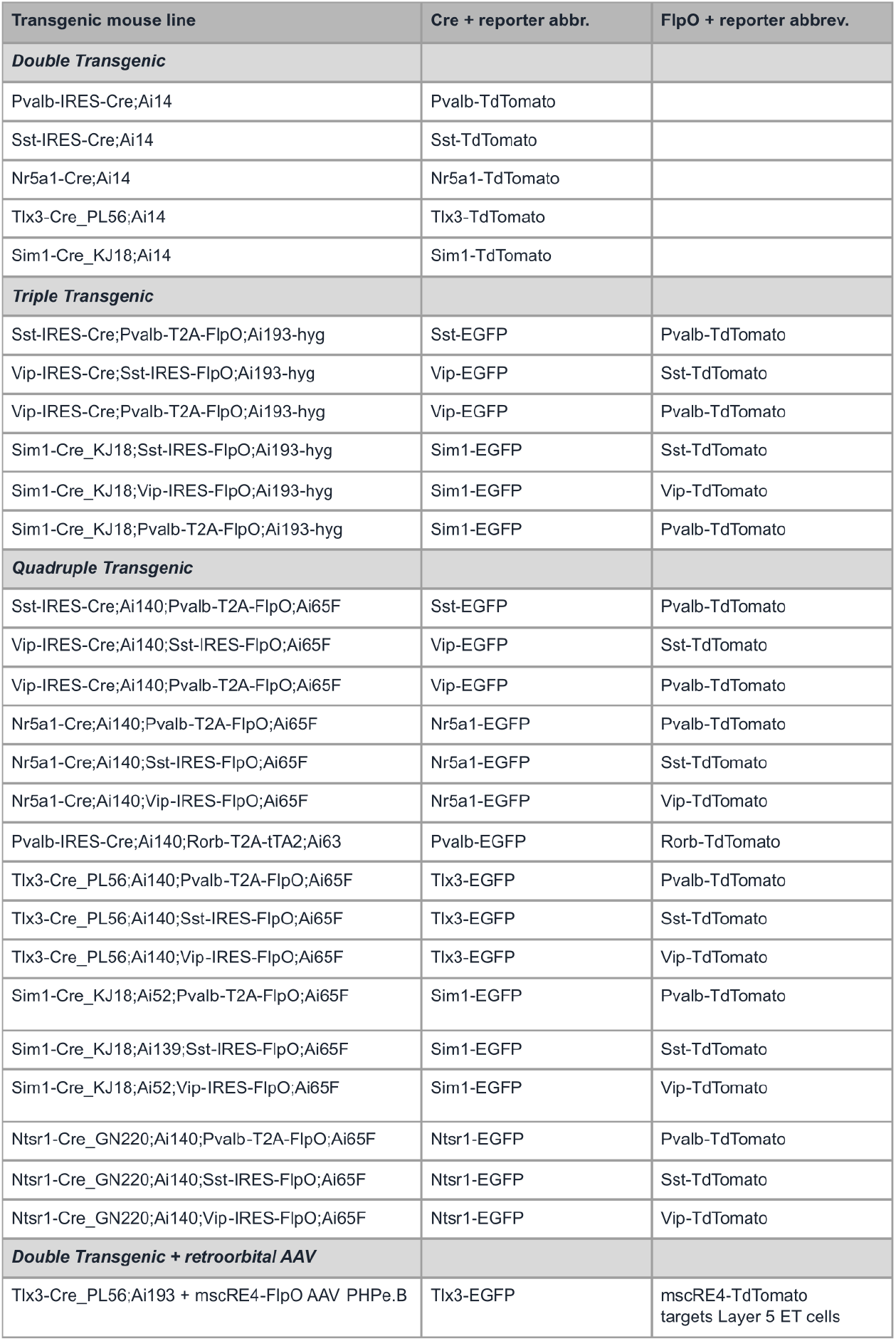
Transgenic Animals. List of transgenic animals used in experiments. Animals were derived from double, triple, or quadruple transgenics. In triple and quadruple transgenics, two subclasses were driven by either Cre or FlpO and expressed either TdTomato or EGFP. In one case we utilized a retroorbital AAV to label L5 ET cells together with L5 IT cells as it was not possible to construct this combination through a triple or quadruple transgenic.

**Table Supplement 2.** ACSF Recipes. Concentrations of each component in ACSF recipes utilized in different stages of our experiments, slice, holding, recording, etc. See Methods for more information on when each ACSF was used.

| Component | ACSF.I | ACSF.IV | ACSF.V | ACSF.VII | ACSF.IX |
| --- | --- | --- | --- | --- | --- |
| Calcium Chloride, dihydrate | 0.5 mM | 2 mM | 1.8 mM | 0.5 mM | 1.3/2.0 mM |
| D-Glucose (Dextrose) | 25 mM | 25 mM | 0 | 25 mM | 25 mM |
| HCl | 98 mM | 0 | 0 | 92 mM | 0 |
| HEPES | 20 mM | 20 mM | 5 mM | 20 mM | 0 |
| Magnesium Chloride | 0 | 0 | 1.0 mM | 0 | 0 |
| Magnesium Sulfate | 10 mM | 2 mM | 0 | 10 mM | 1.0/2.0 mM |
| Monosodium Phosphate | 1.25 mM | 1.25 mM | 0 | 1.2 mM | 1.2 mM |
| Myo inositol | 3 mM | 3 mM | 0 | 0 | 0 |
| N-acetylcysteine | 12 mM | 12.3 mM | 0 | 0 | 0 |
| N-methyl-d-glucamine | 96 mM | 0 | 0 | 92 mM | 0 |
| Potassium Chloride | 2.5 mM | 2.5 mM | 5.4 mM | 2.5 mM | 3mM |
| Sodium Bicarbonate | 25 mM | 25 mM | 0 | 30 mM | 18 mM |
| Sodium Chloride | 0 | 94 mM | 135 mM | 0 | 126 mM |
| Sodium L-Ascorbate | 5 mM | 5 mM | 0 | 5 mM | 0.16 mM |
| Sodium Pyruvate | 3 mM | 3 mM | 0 | 3 mM | 0 |
| Taurine | 0.01 mM | 0.01 mM | 0 | 0 | 0 |
| Thiourea | 2 mM | 2 mM | 0 | 2 mM | 0 |

**Table Supplement 3.** Quality Control. Quality control stages for data processing.

| QC Metric | Analysis level of data being evaluated | Voltage Clamp | Current Clamp |
| --- | --- | --- | --- |
| Holding current | Postsynaptic stimulus trial | $\pm 800$ pA | $\pm 800$ pA |
| Holding potential | Postsynaptic stimulus trial |  | -45 mV to -85 mV |
| Baseline noise | Postsynaptic stimulus trial | < 200 pA | < 5 mV |
| Presynaptic spike detected | Presynaptic recording in response to 1 stimulus pulse | ✓ | ✓ |
| Extra spikes | Presynaptic recording in response to 1 stimulus pulse | None within $\pm 8$ ms | None within $\pm 8$ ms |
| Standard deviation of postsynaptic response<br>This checks for a relatively stable recording | Postsynaptic recording in response to 1 stimulus pulse | < 15 pA | < 1.5 mV |
| Max postsynaptic amplitude<br>This checks for artifacts in the response | Postsynaptic recording in response to 1 stimulus pulse | < 500 pA | < 10 mV |
| Max postsynaptic potential | Postsynaptic recording in response to 1 stimulus pulse |  | -40 mV |
| Excitatory connection | Postsynaptic recording in response to one presynaptic spike | Membrane potential between -45 mV and -85 mV |  |
| Inhibitory connection | Postsynaptic recording in response to one presynaptic spike | Membrane potential between -45 mV and -60 mV |  |

**Table Supplement 4.** Morphology Annotations. Morphological annotations assigned to recorded cells filled with biocytin and imaged at 63x resolution.

**Table Supplement 3 Quality Control.** Quality control stages for data processing.
| Morphology annotation | Description |
| --- | --- |
| ***Dendrite type*** |  |
| NEI (Not Enough Information) | Either no biocytin filled cell or poor health precludes definition of dendrite type |
| Aspiny | No visible spines, non-pyramidal morphology |
| Sparsely spiny | Visible spines, non-pyramidal morphology |
| Spiny | Visible spines, pyramidal or stellate morphology |
| ***Axon*** |  |
| Truncation | Obvious cut axon or retraction bulb near surface of the slice that measured $\leq 100 \mu\text{m}$ |
| Borderline | Axon measured between 100-200 $\mu\text{m}$ |
| Intact | Traceable axon measured $> 200 \mu\text{m}$ |
| Unclear | Axon untraceable but not obviously truncated, likely due to poor health or biocytin fill |
| Truncation distance | Length of axon measured as the straight line distance from origin to end. Only measured in "truncated" or "borderline" axons |
| ***Apical dendrite*** |  |
| Truncated | Does not reach Layer 1 |
| Intact | Reaches Layer 1 |
| Truncation distance | Length of largest apical branch measured as the straight line distance from soma to end of dendrite. Only measured in "truncated" dendrites |

